# Calorie Restriction Induces Degeneration of Neurons with Mitochondrial DNA Depletion by Altering ER-Mitochondria Calcium Transfer

**DOI:** 10.1101/2024.06.14.599123

**Authors:** Lingyan Zhou, Feixiang Bao, Jiajun Zheng, Yingzhe Ding, Jiahui Xiao, Jian Zhang, Yongpeng Qin, Liang Yang, Yi Wu, Qi Meng, Manjiao Lu, Qi Long, Lingli Hu, Haitao Wang, shijuan Huang, Gong Chen, Xingguo Liu

## Abstract

Mitochondrial DNA (mtDNA) mutations and/or depletion are implicated in epilepsy and many neurodegenerative diseases. However, systematic investigation into how mtDNA alterations relate to epilepsy and neural degeneration is needed. Here, we established a mouse model where in mtDNA depletion induced by Herpes Simplex Virus Type 1 (HSV-1) protein-UL12.5 in the brain led to an epileptic phenotype characterized by abnormal electroencephalography (EEG) patterns and increased neural excitability in hippocampus. We also found that UL12.5 mediated mtDNA depletion in neurons *in vitro* (rho^-^) causes epilepsy–like abnormal EEG. Caloric restriction (CR) is a strategy proven to reduce epileptic activity, however CR mimetic 2-deoxy-D-glucose (2-DG), induced degeneration in mtDNA depleted neurons. Mechanistically, mtDNA depletion increased mitochondria-endoplasmic reticulum (ER) contacts, facilitating CR-induced mitochondrial calcium overload. Rho^-^ neurons did not show changes in mitochondrial motility or membrane potential. Our study revealed an unexpected axis of mtDNA depletion, ER-mitochondrial contacts, and calcium overload in the rho^-^ neuron model. This is the first description of animal and neuronal models of mitochondrial epilepsy. Our findings with these models suggest that CR may not be a viable clinical intervention in patients with mtDNA depletion.

## 1. Introduction

Neurological manifestations of mitochondrial dysfunction vary, but a common symptom is epilepsy. Compared to epilepsies in other contexts, mitochondrial epilepsies occur more frequently in the posterior quadrant and occipital lobe, and are more likely to present with non-convulsive status epilepticus which may last months and be more resistant to treatment.[1] The major genetic cause of mitochondrial epilepsy is mtDNA mutations.[2] Mitochondrial respiratory chain deficiency has been reported to promote neuronal hyperexcitability by altering the balance of excitation and inhibition networks.[3] However, another study showed that mtDNA mutations do not impair the synaptic activity of mouse neurons.[4] Despite these findings, there is a significant lack of suitable animal models and mechanistic studies to understand the relationship between mtDNA alterations and epilepsy. This gap has limited the progress in developing targeted therapies and interventions for mitochondrial epilepsy.

CR, defined as a reduction in calorie intake below usual ad libitum intake without malnutrition, showed antiepileptic and anticonvulsant effects in El epilepsy mice models.[5] CR was also shown to decrease neurodegeneration in the brain, and to enhance neurogenesis in animal models of Parkinson’s disease (PD) and Huntington disease.[6] A ketogenic (high-fat, low-carbohydrate) diet, rather than strict CR, showed beneficial effects in mitochondrial epilepsies and CR showed no beneficial action in mtDNA mutator mice.[7]

mtDNA depletion has also been assumed to be a cause of neuronal degeneration.[8] mtDNA depletion syndromes (MDS), which are characterized by a variable tissue-specific reduction of mtDNA copy number, have the most prominent and disabling neurodegeneration features.[9] mtDNA depletion leads to multiple mitochondrial dysfunctions, including impaired oxidative phosphorylation (OXPHOS) and energy deficiency, disturbed lipid-membrane composition, fragmented mitochondria, unbalanced mitochondrial proteins, and dissipated mitochondrial membrane potential.[10] Defective axonal mitochondrial transport causes local energy depletion and toxic changes in calcium buffering that trigger synaptic dysfunction and loss.[11] Alteration of mitochondrial transport has been implicated in neurons from mtDNA-associated neurological disorders.[12] Yet, what kind of mitochondrial dysfunction may promote neural degeneration in mitochondrial epilepsies remains unknown.

Here we establish a mouse model where mtDNA depletion induced by UL12.5 in brains leads to an epileptic phenotype. We also generate a rho^-^ neuronal model (neurons with little or no mtDNA) by expressing HSV-1 UL12.5, and show that treatment with CR mimetic 2-DG, induces degeneration in rho^-^ neurons, which is dependent on ER-mitochondria calcium transfer dysfunction rather than mitochondrial dynamics arrest. Our study reveals an unexpected neuronal degenerative pathway using a novel model of mitochondrial epilepsies and suggests that CR may not be a suitable clinical intervention in patients with mtDNA depletion. Importantly, we have established the first mitochondrial epilepsy mouse and neural model, providing a crucial tool for mechanistic research and potential therapeutic development.

## 2. Results and Discussion

### 2.1 Induction of Epileptic Phenotype in Mice through UL12.5-Mediated mtDNA Depletion

The HSV-1 protein UL12.5, which can localize to the mitochondrial matrix, destroys mtDNA in various cell types, including fibroblasts and Hela cells.[13] To investigate whether reducing mtDNA in the mouse brain could induce epilepsy, we administered PHP.eB-UL12.5 via tail vein injection (Figure 1A-C; Figure S1A, B, Supporting Information). The PHP.eB serotype facilitates virus-specific infection of the brain[14], leading to a decrease in brain mtDNA and mtRNA, without significantly affecting the liver (Figure 1D; Figure S1C, Supporting Information). Further analysis by western blot revealed that UL12.5 significantly reduced the expression of mtDNA-encoded protein MTCOX3, but not nuclear-encoded mitochondrial protein ATP5A (Figure 1E, F).

**Figure 1.**
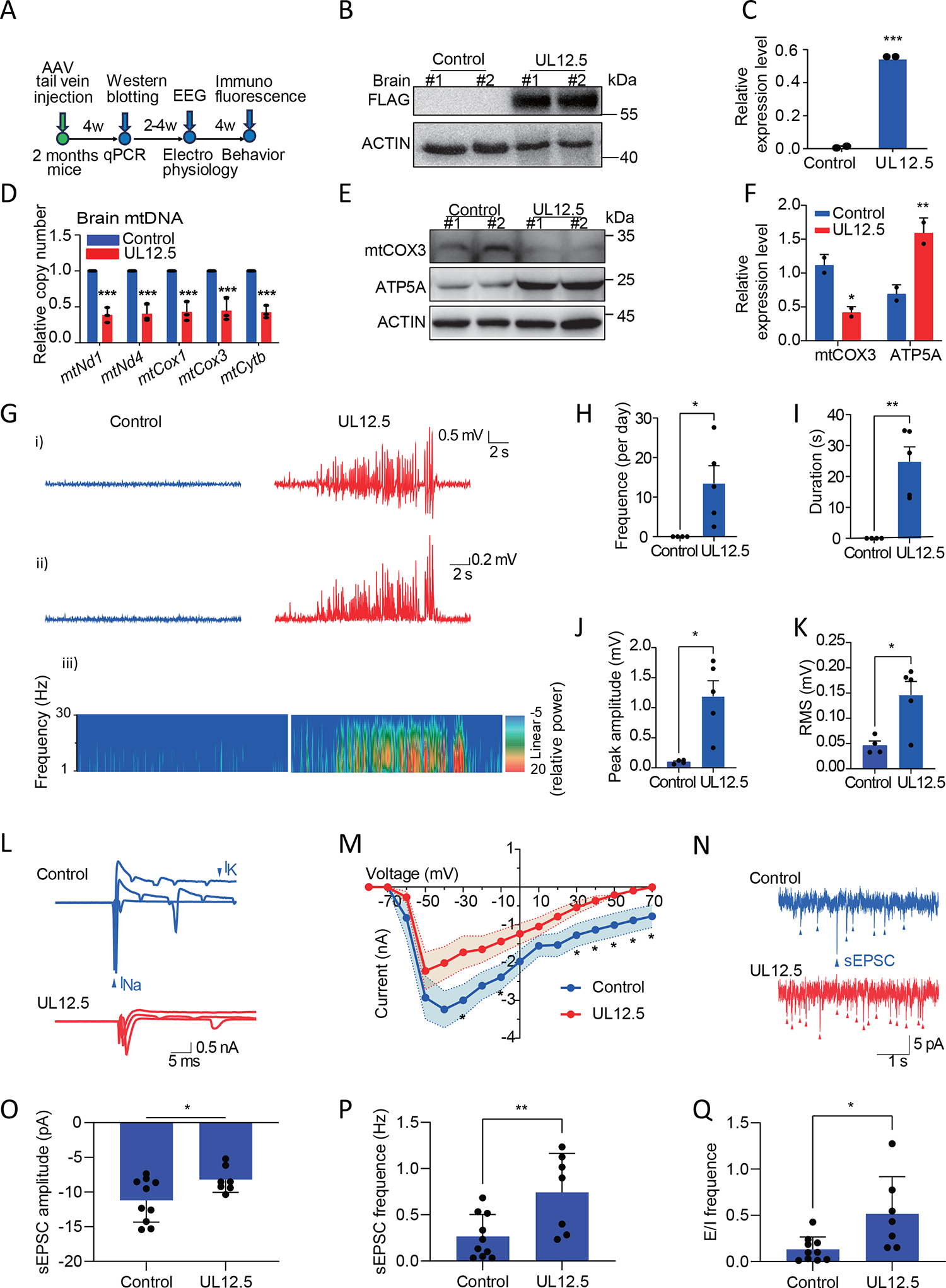
mtDNA depletion in mouse brain induced by UL12.5 resulting in epileptic phenotype. A) Timeline of the mouse model: An AAV injection was administered to deliver 3FLAG or UL12.5-3FLAG AAV at the age of two months, followed by a period of six to eight weeks during which seizures developed in the UL12.5-induced mice. B, C) The Western blot analysis shows overexpression of UL12.5 in mouse brains (B). Quantification of the UL12.5 expression (C). Error bars = SD (n = 2 mice; ***P < 0.001). D) Quantification of the relative mtDNA copy number (*mtNd1*, *mtNd4*, *mtCox1*, *mtCox3*, *mtCytb*) in control and UL12.5 expressing mouse brains by qPCR. Error bars = SD (n = 3; ***p < 0.001). E, F) Western blot analysis of mtDNA-encoded protein (mtCOX3) and nuclear DNA-encoded mitochondrial protein (ATP5A) (E). Quantification of the mtCOX3 and ATP5A expression (F). Error bars = SD (n = 2; *P < 0.05, **P < 0.01). G) Representative EEG and power spectra of the hippocampus from control (left) and UL12.5 mice (right). Root mean square (RMS) of the EEG from control and UL12.5 mice, Spectrograms corresponding to EEG signals were obtained from short-term Fourier transform (STFT). H-K) Quantification of the frequency per day, mean duration, epilepsy peak-amplitude and RMS of EEG. Error bars =SEM (n=4 mice for control, n =5 mice for UL12.5; *P < 0.05, **P < 0.01). L) Representative currents of voltage-dependent sodium and potassium channels in the hippocampus from control and UL12.5 mice. M) The peak amplitude of voltage-activated sodium channel currents (I(Na)) in the hippocampus from control and UL12.5 mice. Error bars: SEM (n =13 neurons from 3 mice; *P < 0.05) N) Representative traces of sEPSCs in hippocampal excitatory neurons from control (blue) and UL12.5 mice (red). O) The amplitude of sEPSCs in hippocampal excitatory neurons from control and UL12.5 mice. Error bars: SD (n =10 neurons from 3 mice for control; n =7 neurons from 3 mice form UL12.5; *P <0.05) P) The frequency of sEPSCs in hippocampal excitatory neurons from control and UL12.5 mice. Error bars: SD (n =10 neurons from 3 mice for control; n =7 neurons from 3 mice for UL12.5; **P <0.01). Q) The excitatory-to-inhibitory (E/I) frequency in hippocampal excitatory neurons from control and UL12.5 mice. Error bars =SD (n =10 neurons from 3 mice for control; n =7 neurons from 3 mice for UL12.5; *P < 0.05).

EEG signals of UL12.5 mice were recorded and analyzed for notable changes in daily frequency, average duration, seizure peak amplitude, and root mean square (RMS). A significant increase in daily mean frequency from 0 to 13.5 Hz was observed in UL2.5 mice, suggesting the presence of intermittent epileptiform abnormal discharges in the hippocampus (Figure 1G-K). Patch clamp whole-cell recording of isolated mouse hippocampal slices revealed a significant reduction in the currents of voltage-dependent sodium and potassium channels and the amplitude of evoked action potentials in UL12.5 mice (Figure 1L; Figure S2A, Supporting Information). Specifically, the amplitude of evoked sodium currents decreased in the hippocampus of UL12.5 mice (Figure 1M; Figure S2B, Supporting Information), which is consistent with changes in the amplitude of excitatory postsynaptic currents (sEPSC) (Figure 1N, O). Interestingly, the frequency of sEPSC was significantly increased, leading to a rise in the E/I ratio (Figure 1P, Q; Figure S2C-E, Supporting Information), which overall reflects a state of hyperexcitability. These results suggest that mtDNA depletion by UL12.5 leads to an epileptic phenotype in mice brain.

Transcriptomic analysis of the brains of UL12.5 mice revealed significant changes in genes related to mtRNA, glutamatergic synapses, apoptosis, axon, and calcium ion homeostasis (Figure S3, Supporting Information). These findings prompted further investigation into the neurons of UL12.5 mice. We evaluated the neurological function of mice using the rotarod test, which showed that a reduction in brain mtDNA significantly decreased the time latency to fall from the rod (Figure 2A). Furthermore, we performed immunofluorescence for NeuN and GFAP, revealing an increase in GFAP-positive glial cells and a decrease in NeuN-positive neuronal cells in the cortex (Figure 2B-D) and the hippocampus (Figure 2E-H).

**Figure 2.**
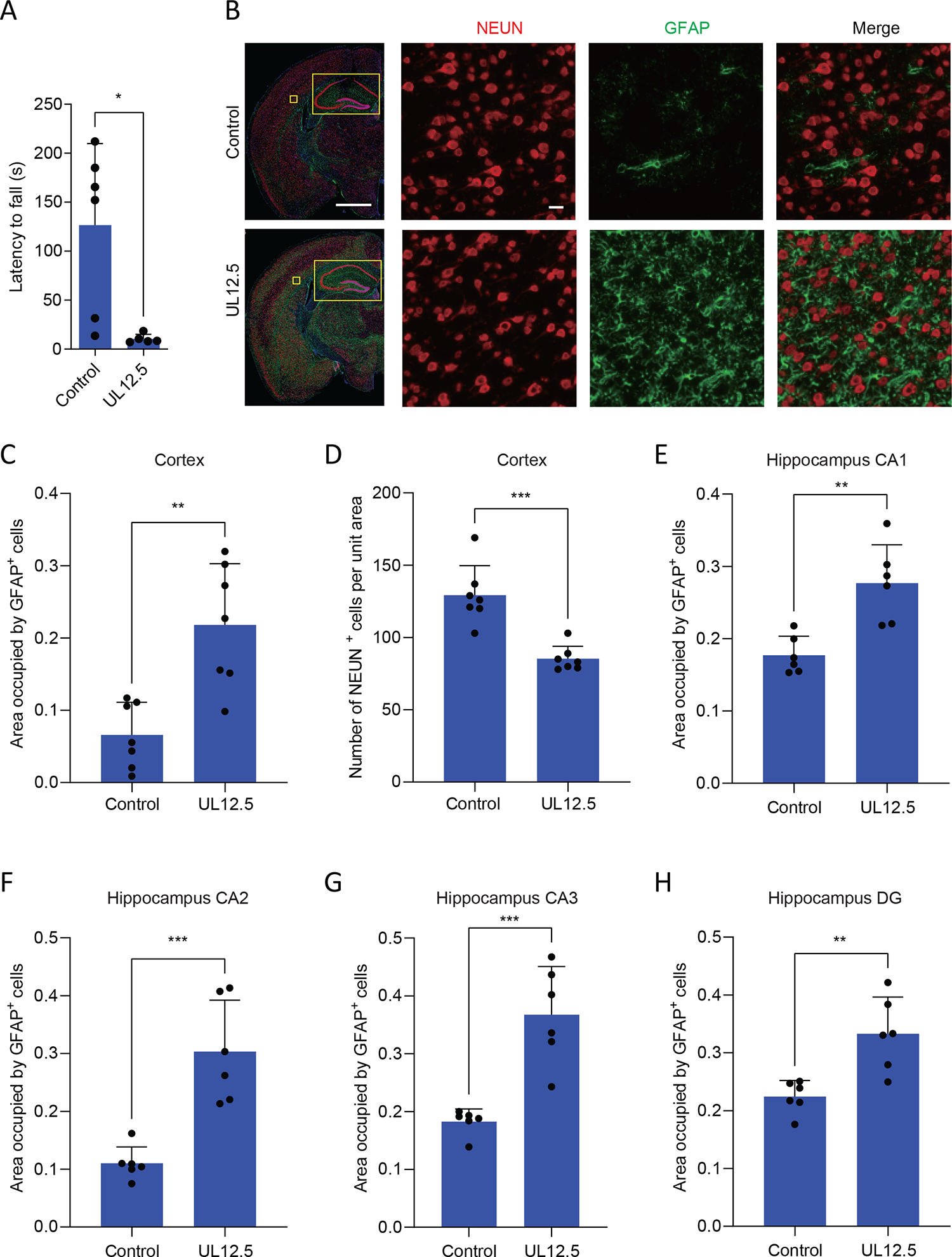
Depletion of mtDNA in the brain leads to cortical and hippocampal neuronal degeneration. A) The latency to fall time in rotarod tests of 3FLAG and UL12.5-3FLAG mice 10-12 weeks after AAV injection. Error bar: SD (n ≥ 5 mice; *p < 0.05). B-D) Representative images of UL12.5 expressing brain with markers of glia (GFAP) and neuron (NEUN) (B). Scale bars: 1 mm (overview) and 100 μm (insert). Quantification of GFAP and NEUN stained cells in cortex and hippocampus (C, D). Error bars: SD (n = 7 mice; **P < 0.01, ***P < 0.001). E-H) Quantification of GFAP-positive glia in hippocampal regions: Cornu Ammonis 1 (CA1), Cornu Ammonis 2 (CA2), Cornu Ammonis 3 (CA3), and Dentate Gyrus (DG). Error bars: SD (n = 6 mice; **P < 0.01, ***P < 0.001).

### 2.2 UL12.5 Expression Causes rho^-^ Neuron Degeneration Following 2-Deoxyglucose Treatment

We tested whether UL12.5 could also deplete neural mtDNA to generate rho^-^ neurons. We expressed UL12.5 fused with fluorescent protein (DsRed) in human induced pluripotent stem cell-derived neurons. Immunostaining showed UL12.5 localized in mitochondria (Figure S4A, Supporting Information). To investigate the effect of UL12.5 on neural mtDNA, we labeled mtDNA and mitochondrial nucleoids with anti-DNA and anti-TFAM antibodies, respectively. UL12.5 expression significantly decreased mtDNA mass and also destroyed nucleoid structure (Figure 3A, B; Figure S4B, Supporting Information). Similar results were obtained by detecting mtDNA copy number using qPCR (Figure 3C). Transcription and translation of mtDNA in rho^-^ neurons were also abolished (Figure S4C, Supporting Information), and mtDNA encoded protein MTATP8, but not nuclear encoded mitochondrial protein ATP5A, was lost in neurons expressing UL12.5 (Figure 3D). Thus, UL12.5 expression decreased neural mtDNA and could be used to construct mtDNA depletion rho^-^ neuron model.

**Figure 3.**
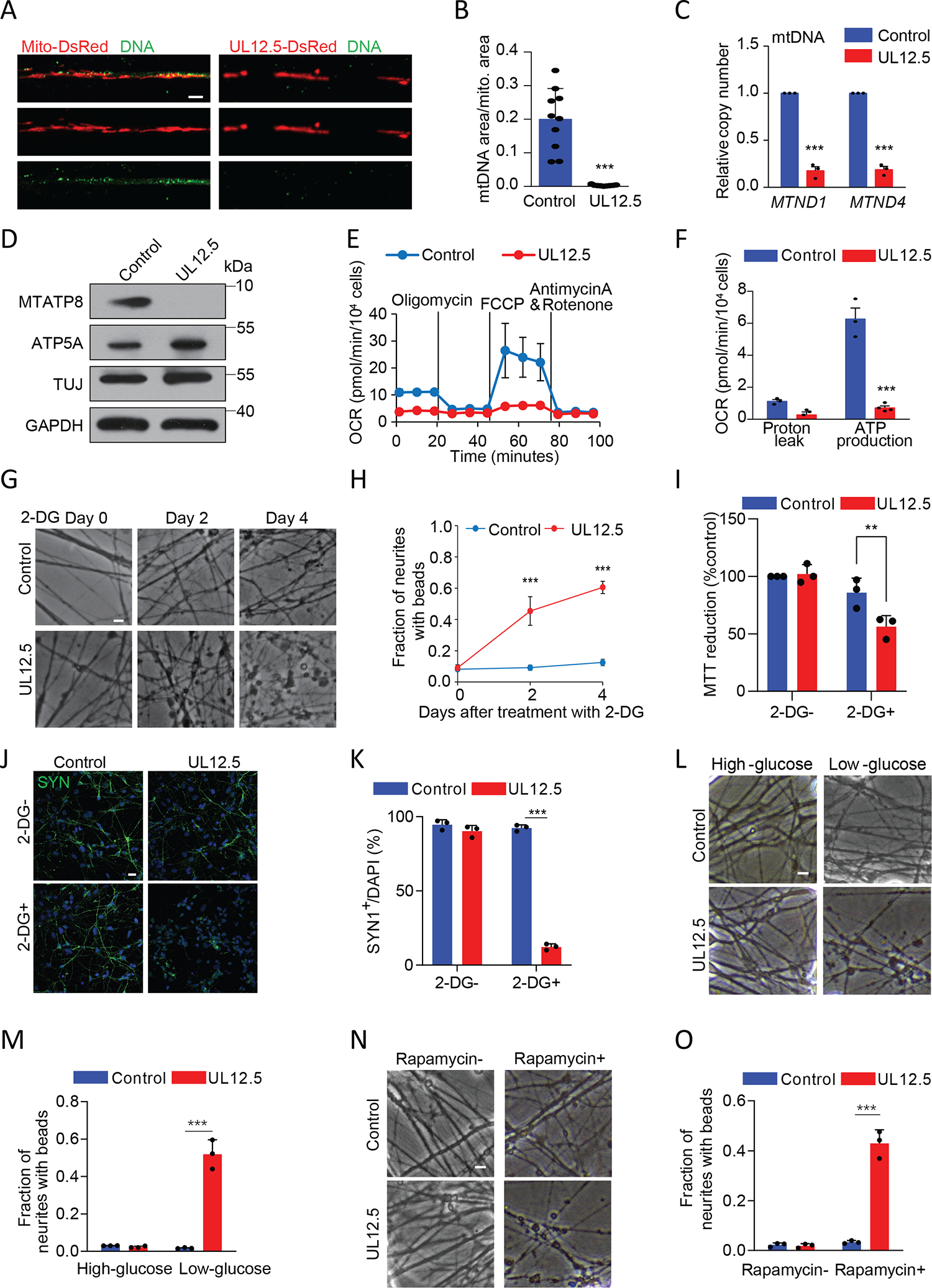
CR induces degeneration of UL12.5 expressed rho^-^ neurons. A) Representative images of a fixed UL12.5-DsRed (red) expressing neuron with mitochondrial marker protein TOM20 (green). Scale bar: 10 μm. B) Quantification of the ratio of mtDNA area to mitochondrial area. Error bars: SD (n = 10 neurons; ***P < 0.001). C) qPCR was used to detect the relative mtDNA copy number (*MTND1*, *MTND4*) in control or UL12.5-DsRed expressing neurons. Error bars: SEM (n = 3; ***p < 0.001). D) Western blot analysis of mtDNA-encoded protein (MTATP8) and nuclear DNA-encoded mitochondrial protein (ATP5A). E, F) Mitochondrial respiration was abrogated in UL12.5 expressing neurons. Neurons were infected with mito-DsRed or UL12.5-DsRed before OCR measurement. Oligomycin (inhibiting ATP synthesis), FCCP (an uncoupler inducing maximum respiration), and antimycin A & rotenone (inhibiting total mitochondrial respiration) were added sequentially (E). (F) Proton leak (subtraction of the antimycin A & rotenone OCR from the oligomycin OCR) and mitochondrial ATP production capacity (subtraction of the oligomycin OCR from the basal OCR). Error bars: SEM (n ≥ 3; ***P < 0.001). G, H) Control or rho^-^ neurons were treated with 2-DG for 0, 2, and 4 days. Representative brightfield images are shown in (G). Scale bar: 25 μm. Quantification of the fraction of neurites with beads (H). Error bars: SD (n = 4; ***p < 0.001). I) Quantification of the MTT reduction in control and rho^-^ neurons. Error bars: SD (n = 3; **p < 0.01). J, K) Representative immunofluorescence images of synapses (SYN1, green) and nuclei (DAPI, blue) (J), scale bar: 10 μm. Quantification of the ratio of SYN1 spots to cells (K). Error bars: SD (n = 3; ***p < 0.001). L, M) Representative brightfield images of control and rho^-^ neurons treated with high-glucose or low glucose for 4 days (L). Scale bar: 25 μm. Quantification of the fraction of neurites with beads (M). Error bars: SD (n = 3; ***p < 0.001). N, O) Representative brightfield images of control and rho^-^ neurons treated with rapamycin for 4 days (N). Scale bar: 25 μm. Quantification of the fraction of neurites with beads (O). Error bars: SD (n = 3; ***p < 0.001).

Decreased expression of mitochondrial encoded electron chain components is expected to lead to metabolic dysfunction. We measured cellular metabolism using the Seahorse Flux Analyzer and found mitochondrial ATP production and maximal respiration were greatly reduced in neurons expressing UL12.5 (Figure 3E, F; Figure S5A, Supporting Information), the analysis of glycolysis indicates that the ability to enhance glycolysis upon inhibition with oligomycin is diminished, suggesting that the maximum glycolytic capacity might have already been achieved (Figure S5B-D, Supporting Information). However, baseline glycolytic activity (glucose - 2-DG) remains unchanged. This observation implies that the rho^-^ neurons maintain adequate glycolytic activity to compensate for the decreased energy production resulting from impaired OXPHOS.

Given that epilepsy is a major phenotypic feature in mtDNA depletion diseases,[15] we asked whether acute mtDNA depletion could cause epilepsy-like pathological changes in neuronal electrophysiology. Expression of UL12.5 led to reductions in both Na^+^ and K^+^ currents and action potential amplitudes (Figure S6, Supporting Information), indicating an epilepsy-like phenotype in our rho^-^ neuron model.

CR is able to reduce epileptic activity by inhibiting the mammalian target of rapamycin (mTOR) signaling pathway.[16] However, CR showed no benefit on the lifespan and healthspan of POLG mitochondrial mutator mice, suggesting that mtDNA dysfunction may interfere with the beneficial effects of CR.^[7a]^ We administered CR mimetic 2-DG (a competitive inhibitor of glucose metabolism) to rho^-^ neurons. Successful induction of a CR phenotype by 2-DG was validated through qPCR, which detected upregulation in the expression of *SIRT1*, *PPARGC1A*, *mTOR*, *PRKAB1*, and *PRKAA2* (Figure S7A, Supporting Information). Surprisingly, 2-DG treatment induced the degeneration of rho^-^ neurons (Figure 3G, H). The 2-DG induced degeneration was further validated by MTT assay and cell viability studies (Figure 3I; Figure S7D, Supporting Information). Immunofluorescence staining for synapsin I (SYN1), a marker of mature neuron synapses, showed a marked reduction in SYN1-positive cells among rho^-^ neurons (Figure 3J, K).

To explore whether alternative CR strategies could induce similar neurodegenerative effects as 2-DG, low-glucose culture conditions and rapamycin treatment were utilized. The results indicated that both approaches also resulted in significant neurodegeneration in rho^-^ neurons (Figure 3L-O; Figure S7B, C and E, F, Supporting Information).

### 2.3 2-DG Induced Rho^-^ Neurons Degenerate Independent of Mitochondrial Transport or **Δψ**_m_

Many neurodegenerative diseases that are associated with decreased mtDNA, including PD and amyotrophic lateral sclerosis (ALS), are also characterized by axonal mitochondrial transport disruption.[12, 17] We interrogated whether disrupted mitochondrial transport following mtDNA depletion underlies increased sensitivity to 2-DG (Figure 4A). We labeled mitochondria with mito-DsRed or UL12.5-DsRed and measured mitochondrial movement in neurites using kymograph analysis. Unexpectedly, neither mtDNA depletion nor 2-DG treatment (12 h), nor the combination had a detectable effect on mitochondrial transport (Figures 4B-E; Figure S8, Supporting Information). These results exclude the dysfunction of mitochondrial movement as degenerative mechanisms of rho^-^ neurons induced by 2-DG.

**Figure 4.**
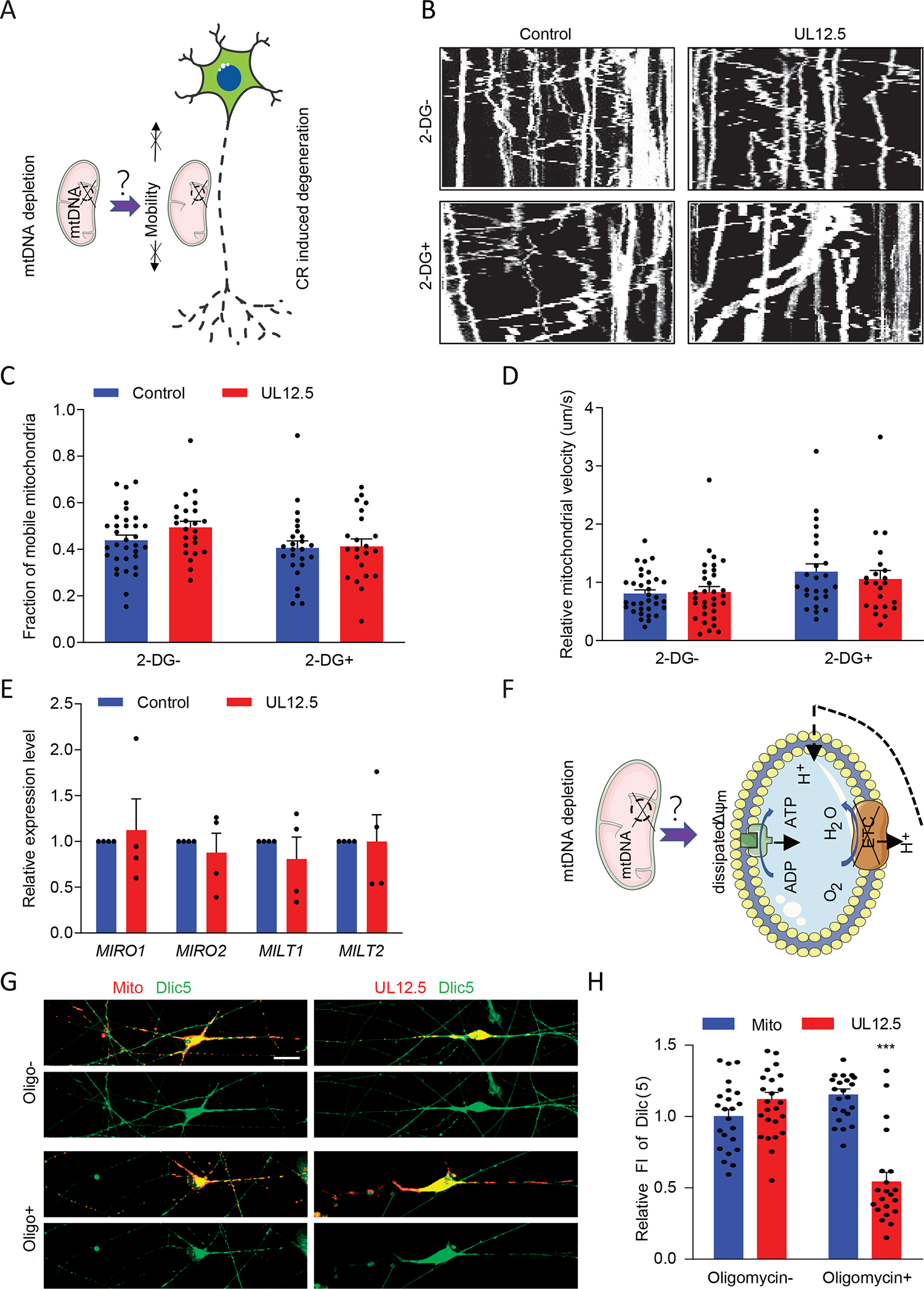
Rho^-^ neurons degenerate after CR treatment independently of mitochondrial transport or Δψ_m_. A) Schematic of the hypothesis: mitochondrial transport disruption contributes to CR induced degeneration. B,C) 2-DG does not affect mitochondrial motility in rho^-^ neurons. Representative mitochondrial kymographs recorded from control and rho^-^ neurons with or without 2-DG treatment for 12 h are shown in (B). The percentage of motile mitochondria is shown in (C). Error bars: SEM (n ≥ 22 mitochondria). D) Analysis of the velocities of mobile mitochondria in each direction, (mitochondrial velocity was normalized to the control without 2-DG). Error bars: SEM (n≥22 mitochondria). E) qPCR analysis of mitochondrial dynamics related genes (*MIRO1*, *MIRO2*, *MILT1* and *MILT2*) expression in control and rho^-^ neurons. Error bars: SEM (n = 4). F) Schematic of the hypothesis: mtDNA depletion contributes to Δψ_m_ dissipation. G, H) UL12.5-DsRed or Mito-DsRed expressing neurons treated with or without 1 μg/mL oligomycin for 1 hour were loaded with Dilc (5) for in the last 30 min. Representative confocal microscopy images (G). Scale bar: 10 μm. Quantification of the Dilc (5) fluorescence intensity is shown in (H). Fluorescence intensity was normalized to the control without oligomycin. Error bars: SEM (n≥22 neurons; ***p < 0.001).

Since mitochondrial membrane potential (Δψ_m_) was shown to decrease in rho0 Hela cells and fibroblasts,^[13b]^ we investigated whether dissipated Δψ_m_ in rho^-^ neurons is responsible for degeneration in response to 2-DG (Figure 4F). No significant difference in Δψ_m_ was observed in rho^-^ neurons, using the membrane-potential sensitive, far-red fluorescent dye 1,1’,3,3,3’,3’-hexamethylindodicarbo-cyanine iodide (Dilc (5)) (Figure 4G, H). Considering that the H^+^-ATP synthase can function in reverse, acting as an ATP hydrolase for maintaining the proton motive force,[18] we hypothesized that Δψ_m_ was maintained via this mechanism in rho^-^ neurons. Indeed, Δψ_m_ was substantially lost in rho^-^ neurons treated with oligomycin (a specific F_1_-F_0_ ATP synthase inhibitor) (Figure 4G, H). Similar results were recently found in NADH dehydrogenase [ubiquinone] iron-sulfur protein 2 (NDUFS2)-knockout dopaminergic neurons with disrupted functional mitochondrial complex I, wherein blocking the adenine nucleotide transporter caused mitochondrial potential to collapse.[19] Collectively, these results indicated that mitochondrial mobility and Δψ_m_ are preserved in rho^-^ neurons.

### 2.4 CR Induces Degeneration of rho^-^ Neurons through an ER-Mitochondria Calcium Transfer Mechanism

We previously reported that Δψ_m_-dependent, tight associations between mitochondria and fragmented ER are involved in calcium dysfunction during neural degeneration.[20] Mitochondrial calcium signaling dysfunction may also contribute to CR-induced degeneration (Figure 5A). This led us to examine ER morphology in mtDNA depleted neurites using ER-targeted DsRed. Neuritic ER fragmentation was increased in rho^-^ neurites compared to control, indicative of an impaired ER homeostasis (Figure 5B, C). We used a contact reporter system based on split-GFP (ER-GFP(1-10) and mito-GFP11)[21] to measure ER-mitochondrial contacts in neurons. By both flow cytometry and confocal imaging we found higher split GFP signal in rho^-^ neurons than in controls (Figure 5D, E; Figure S9A, B, Supporting Information).

**Figure 5.**
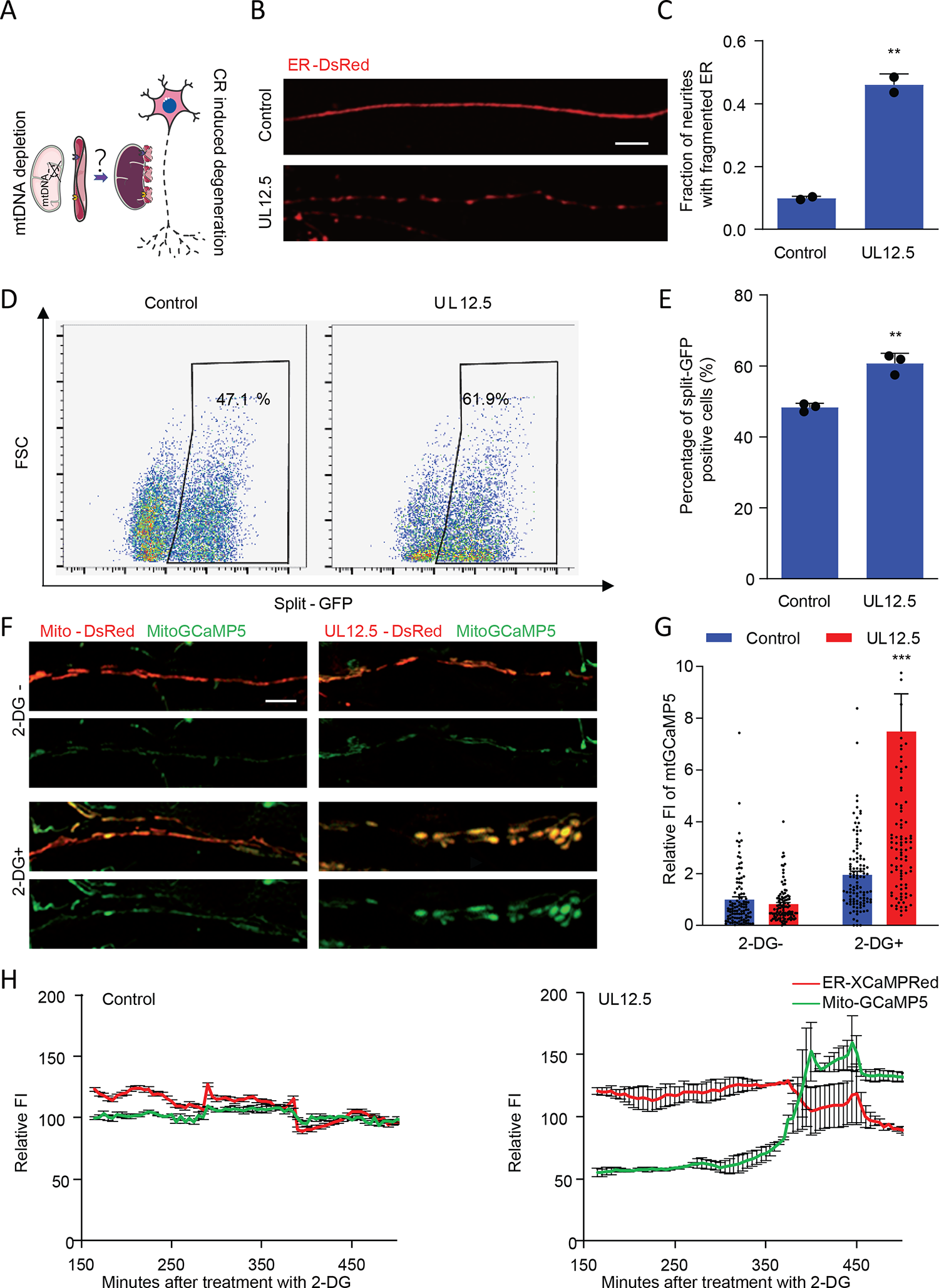
CR-induced degeneration of rho^-^ neurons depends on ER-mitochondria contacts. A) Schematic of the hypothesis: calcium signal dysfunction contributes to CR induced degeneration. B, C) Neuritic ER fragmentation in rho^-^ neurons. Neuritic ER was labeled with ER-DsRed and representative confocal microscopy images are shown (B). Scale bar: 10 μm. Quantification of the fraction of neurites with fragmented ER (C). Error bars: SD (n = 2; **p < 0.01). D, E) Neurons expressing split-GFP to label mitochondria-ER contacts show that the contact of mitochondria-ER increases in rho^-^ neurons. Representative flow cytometry plots are shown (D). Quantification of the fraction of split-GFP positive cells (E). Error bars: SD (n = 3; ** p < 0.01). F, G) Control and rho^-^ neurons expressing mito-GCaMP5 were treated with 2-DG for 2 days. Representative confocal microscopy images (F). Scale bars: 10 μm. Quantification of the mito-GCaMP5 fluorescence intensity is shown in (G). Fluorescent intensity was normalized to the control neurons without 2-DG. Error bars: SEM (n ≥ 100 neurons; *** p < 0.001). H) Control and rho^-^ neurons expressing mitochondrial GCaMP5 and ER XCaMPRed were monitored following treatment with 2-DG. The fluorescence intensities of GCaMP5 and XCaMPRed were normalized to 100 at time=0. Error bars: SEM (n ≥ 3).

Considering that dysregulated mitochondria-ER contacts are associated with calcium dysfunction[22] and that CR enhances brain mitochondrial calcium retention capacity,[23] we reasoned that calcium dysfunction could factor in CR-induced rho^-^ neuron degeneration. We checked whether 2-DG affects cytosolic calcium levels in rho^-^ neurons. Fluo-4 acetoxymethyl (Fluo-4) staining showed that 2-DG triggered an increase in intracellular calcium (Supplementary Figure S9C, D, Supporting Information). Importantly, 2-DG also increased mitochondrial calcium in rho^-^ neurons, as measured by mito-GCaMP5 fluorescence (Figure 5F, G). To monitor calcium transfer from the ER to the mitochondria, we simultaneously expressed the ER calcium probe ER-XCaMPRed and the mitochondrial calcium probe mito-GCaMP5 in neurons. Following 2-DG treatment, live cell imaging demonstrated an increase in fluorescence intensity for mito-GCaMP5 and a concomitant decrease for ER-XCaMPRed (Figure 5H; Figure S9E-G, Supporting Information). This suggests that calcium transfer between the ER and mitochondria contributes to the neurodegenerative effects induced by 2-DG.

Increases in cytosolic calcium concentration can be caused by calcium release from intracellular stores, mainly ER, or by calcium influx from the extracellular space.[24] To investigate the source of calcium for the CR-induced degeneration of rho^-^ neurons, we used extracellular calcium chelation by EGTA or intracellular calcium chelation by BAPTA-AM together with 2-DG treatment. BAPTA-AM rather than EGTA clearly prevented the beads formation of rho^-^ neuron (Figure 6A, B), indicating an intracellular source of calcium rather than extracellular calcium influx participates in 2-DG induced rho^-^ neuronal degeneration. Transcriptome profiling analysis revealed significant alterations in the expression of genes associated with mitochondrial and ER interactions, particularly affecting calcium channels like inositol 1,4,5-trisphosphate receptors (IP3R), ryanodine receptors (RyR), voltage-dependent anion channel (VDAC), and MCU (Figure S10, Supporting Information). Calcium-selective channels, known as IP3Rs and RyR, regulate ER calcium release and ER-mitochondria calcium transfer.[25] Increased neuronal IP3R- and RyR-mediated calcium release has been observed in degenerative diseases.[26] To determine whether calcium transfer from ER to mitochondria is involved in this process, we used ER calcium channel inhibitors Dantrolen (Dan, RyR antagonist) and Xestospongin C (XesC, IP3R antagonist). Both Dan and XesC prevented the degeneration of rho^-^ neurons induced by 2-DG (Figure 6A, B). Given the limited specificity of Dan and XesC, we explored more specific antagonists, namely Ryanodine for RyR and 2-APB for IP3R. The results showed that both Ryanodine and 2-APB also effectively inhibited the neurodegeneration induced by 2-DG (Figure 6C; Figure S11, Supporting Information). Furthermore, we conducted shRNA-mediated knockdown of *IP3R3* and *RYR1*, which were up-regulated in rho^-^ neurons (Figure S12, Supporting Information). The knockdown of either *IP3R3* or *RYR1* was also able to block neurodegeneration induced by 2-DG in rho^-^ neurons (Figure 6D, E). Taken together, CR induced degeneration of rho^-^ neurons is dependent on increased ER-mitochondrial calcium transfer.

**Figure 6.**
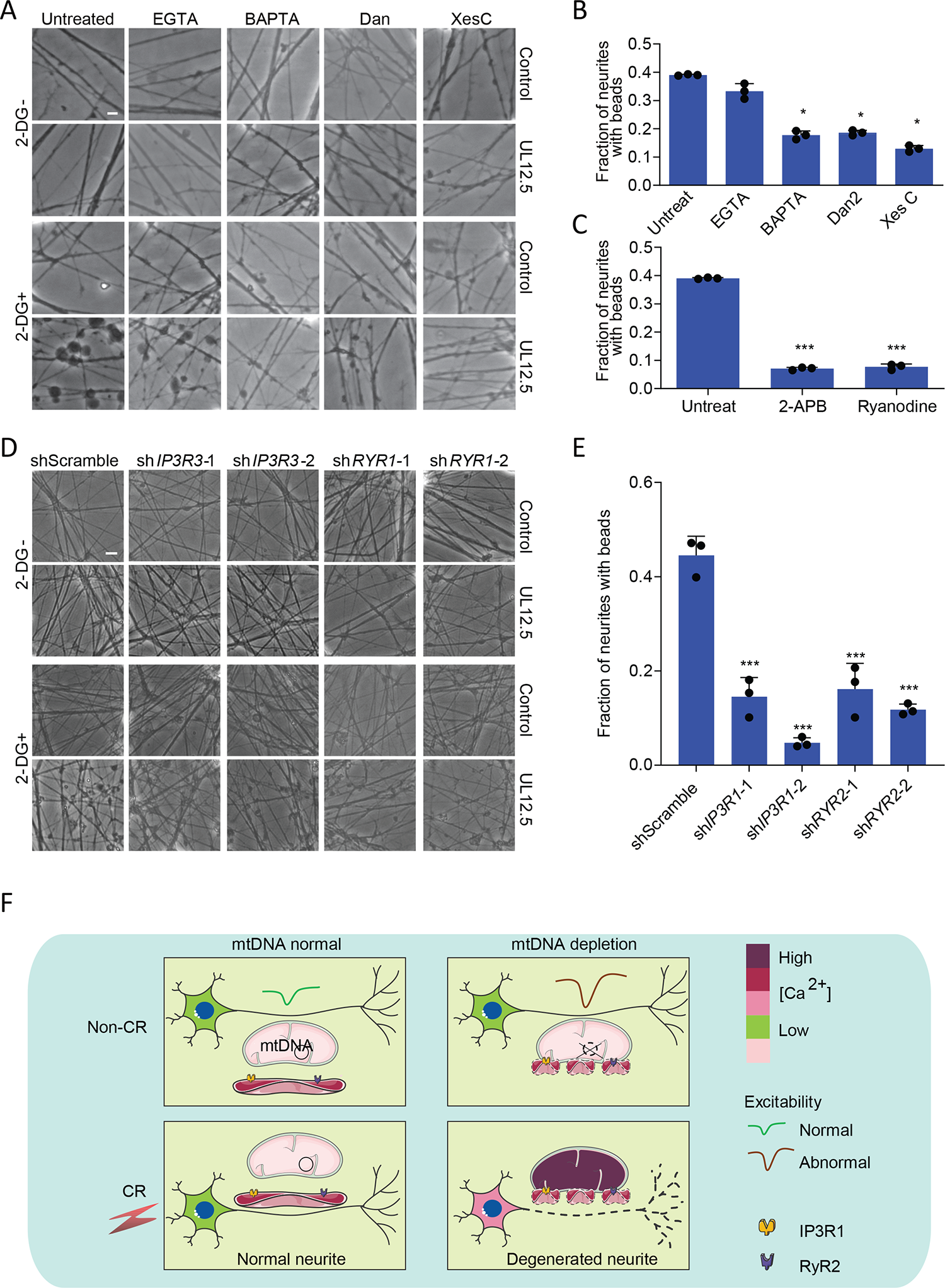
The IP3R and RyR calcium channels are involved in the degeneration of rho^-^ neurons induced by CR. A, B) Neuritic degeneration induced by 2-DG in rho^-^ neurons was prevented by BAPTA-AM, Dan or XesC. A) Images of control and rho^-^ neurons after 2-DG with or without simultaneous EGTA, BAPTA-AM, Dan or XesC treatment for 2 days. Scale bar: 10 μm. B) Fractions of neurites with beads. Error bars: SD (n = 3; *p < 0.05). C) Neuritic degeneration induced by 2-DG in rho^-^ neurons was prevented by 2-APB or Ryanodine. Fractions of neurites with beads. Error bars: SD (n = 3; ***p < 0.001). D, E) Neuritic degeneration induced by 2-DG in rho^-^ neurons was prevented by *IP3R3* or *RYR1* knock down. D) Representative images of control and rho^-^ neurons after 2-DG with shScramble, sh*IP3R3* or sh*RYR1*. Scale bar: 10 μm. E) Fractions of neurites with beads. Error bars: SD (n = 3; ***p < 0.001). F) Model of 2-DG induced degeneration in rho^-^ neurons.

## 3. Discussion

In this study, we developed the first mitochondrial epilepsy animal model, which exhibits abnormal EEG patterns and increased neural excitability in the hippocampus. We also reported a novel mtDNA depleted, rho^-^ neuronal model in which abnormal electrophysiology, i.e. Na^+^ and K^+^ currents and evoked action potentials are observed, and which may reflect mitochondrial epilepsies caused by mtDNA abnormalities. Epilepsy is the most common clinical presentation of mitochondrial diseases,[27] though neurons with a variety of mtDNA mutations showed normal synaptic activity.[4] Mitochondrial dysfunction induced by rotenone has been reported to cause the loss of inward current.[28] Epilepsy phenotypes may be associated with mtDNA mutated sites and mitochondria-related nuclear gene mutations, which make it difficult to model. Different mtDNA mutations give rise to distinct syndromes and epilepsy phenotypes, including mitochondrial encephalomyopathy with lactic acidosis and stroke-like episodes (MELAS), myoclonic epilepsy and ragged red fibers (MERRF), progressive external ophthalmoplegia (PEO), Leber hereditary optic neuropathy (LHON), Kearns-Sayre syndrome, and Leigh disease.[29] Our mouse model of mitochondrial epilepsy and the mtDNA-depleted rho^-^ neurons provide valuable tools for mechanistic studies of mitochondrial epilepsy and can be useful to evaluate drug candidates for this condition.

CR reduces epileptic activity by inhibiting the mTOR signaling pathway,[16] but has no beneficial action in mice with mtDNA depletion caused by polymerase gamma (POLG) mutation.^[7a]^ Here, we report that CR mimetic 2-DG induces neural degeneration in mtDNA depleted rho^-^ neurons, suggesting that the clinical usage of CR, especially for patients with mitochondrial dysfunction, should be reevaluated due to a risk of neural degeneration.

We report a novel axis involving mtDNA depletion, ER-mitochondrial contacts, and calcium overload, which facilitates ER calcium release and mitochondrial calcium overload (Figure 6F). Mitochondria-ER contact sites (MERCs) control the dynamic distance and cross-talk between ER and mitochondria.[30] MERCs are known to underpin many important cellular functions, including lipid metabolism, calcium homoeostasis, unfolded protein response, ER stress and mitochondrial quality control.[31] We previously reported that MERC-associated mitochondrial calcium overload participates in neuronal degeneration.[20, 32] Mutations of MERC-resident proteins Thioredoxin Related Transmembrane Protein 2 (TMX2) or Serine Active Site Containing 1 (SERAC1) have also been reported to cause epilepsy.[33] Abnormal ER-mitochondria contact seems to be a quite universal process occurring in multiple pathological settings, including nonalcoholic fatty liver disease, PD and ALS.[34] We also found changes in the activity and expression of proteins that mediate ER-mitochondrial calcium signaling (IP3R and RyR) in the rho^-^ neurons. Addressing the role of the mtDNA depletion-MERCs-calcium axis in mitochondrial epilepsies demands future studies.

Here we clarified the relationship between mitochondrial movement and mtDNA depletion in neuronal degeneration. Abnormal mitochondrial movement is an important cause for neural degeneration in many mtDNA alteration-associated degenerative diseases, such as Alzheimer’s disease and PD.[35] However, in our rho^-^ model, mitochondrial movement is normal, showing that mitochondrial movement is not directly affected by the reduction of mtDNA copy number. The lack of energy production by mitochondrial OXPHOS in our rho^-^ neurons suggests that mitochondria might maintain their mobility through a compensatory pathway, analogous to the mechanism of vesicular transport supported by glycolysis-generated energy in axons; recent studies have indicated an enrichment of glycolytic enzymes alongside vesicles in axonal regions.[36] Additionally, anaerobic respiration and the pentose phosphate pathway in the cytosol may also produce ATP,[37] thus potentially participating in mitochondrial movement. Mitochondrial movement is regulated by the cytosolic ADP/ATP ratio and calcium concentration. The AMPK–PAK–myo6–SNPH pathway enables neurons to recruit presynaptic mitochondria at regions of high energy demand.[38] The Miro–Milton complex can enable neurons to efficiently retain mitochondria at the sites with high calcium to buffer calcium,[39] Given the importance of mitochondrial transport to axons, neurons may contain multiple redundant energy supply mechanisms to ensure the movement of mitochondria. Under normal conditions, mitochondrial ATP provides the energy for their movement. Once this pathway is inhibited, alternative energy sources such as glycolysis can take over to maintain mitochondrial movement. We observed that both mitochondrial membrane potential and transport are preserved in rho^-^ neurons. It remains unclear whether mitochondrial transport depends on membrane potential under conditions of mitochondrial ATP depletion.

## 4. Conclusion

It was demonstrated in this study that the depletion of mtDNA from mouse brains through UL12.5 results in abnormal EEG patterns and increased neural excitability in the hippocampus. The expression of UL12.5 destroys neural mtDNA, leading to the generation of rho^-^ neuron cells. Rho^-^ neurons with epilepsy-like electrophysiology may be a good model for mitochondrial epilepsy. Importantly, rho^-^ neurons are more susceptible to neural degeneration caused by CR mimetic 2-DG than neurons with normal mtDNA. Without changing mitochondrial motility or membrane potential, mtDNA depletion leads to increased mitochondria-ER contacts, thus facilitating CR-induced mitochondrial calcium overload in neurons (Fig. 6F).

## 5. Experimental Section Mice

All housing, breeding, and other animal procedures were approved by the Guangzhou Institutes of Biomedicine and Health (GIBH) Ethical Committee and in accordance with GIBH Institutional Animal Care and Use Committee guidelines (Approve no. 2018040). Chronic EEG and whole-cell patch-clamp recordings was approved from the GHM Institute of CNS Regeneration, Jinan University. Wild-type C57BL/6DJ mice aged 2 months were purchased from the BesTest Bio-Tech Co,.Ltd. (Zhuhai, China). Mice were maintained under standard laboratory conditions (12-hour light/dark cycle, room temperature 21° ± 1°C) with free access to water and food and were adapted to the laboratory conditions for at least 1 week.

### AAV tail vein injection

Tail vein injection of either PHP.eB-control or PHP.eB-UL12.5 were performed at 8 weeks of age. Adenoviruses encoding 3FLAG or UL12.5-3FLAG were given to mice at the dose of 1-3 × 10^11^ virus particles by injection into the tail vein. After 4 weeks, livers and brains were collected.

### EEG recordings

EEG recordings were performed in freely moving mice during the chronic phase of epilepsy. In all animals, recordings began between 6-8weeks after AAV tail vein injection. The left hippocampi were implanted with electrodes for chronic EEG recordings. The animals were anesthetized with isoflurane (1−2.5%) during the implantation procedure. Digital EEG activity was monitored daily for up to 3 days during 24-h sample recordings. A digital video camera was used to simultaneously monitor behavior during the EEG recordings. All recordings were carried out at least 3 days after surgery on mice freely moving in the cage. Data was collected by Spike2 8.0.

### Whole-cell patch-clamp recordings

Hippocampal slices were prepared typically 6-10 weeks after virus injection and cut at 300 μm thick horizontal slices with a Leica vibratome (VT-1200S) in ice-cold cutting solution (containing 75 mM sucrose, 85 mM NaCl, 2.5 mM KCl, 0.5 mM CaCl₂, 4 mM MgCl₂, 24 mM NaHCO₃, 1.25 mM NaH₂PO₄ and 25 mM glucose). Slices were incubated in NMDG-ACSF (containing 93 mM NMDG, 2.5 mM KCl, 1.2 mM NaH₂PO₄, 30 mM NaHCO₃, 20 mM HEPES, 25 mM glucose, 5 mM sodium ascorbate, 2 mM Thiourea, 3 mM sodium pyruvate, 10 mM MgSO₄⋅7H₂O, 0.5 mM CaCl₂, pH 7.3 adjusted with HCl, 300–310 mOsm/L), continuously bubbled with 95 % O₂ and 5 % CO₂, first at 34 ◦C for 30 min, and then at room temperature. Whole-cell recordings were performed using Multiclamp 700B patch-clamp amplifier (Molecular Devices, Palo Alto, CA), and the slices were maintained in artificial cerebral spinal fluid (ACSF) containing 126 mM NaCl, 2.5 mM KCl, 1.25 mM NaH₂PO₄, 26 mM NaHCO₃, 2 mM MgCl₂, 2 mM CaCl₂ and 10 mM glucose. The pH of bath solution was adjusted to 7.3 with NaOH, and osmolarity at 310–320 mOsm/L. Patch pipettes were pulled from borosilicate glass (∼5− 8 MΩ) and filled with a pipette solution consisting of 126 mM K-Gluconate, 4 mM KCl, 10 mM HEPES, 4 mM Mg₂ATP, 0.3 mM Na₂GTP, 10 mM Phospho-Creatnine (pH 7.3 adjusted with KOH, 290 mOsm/L). Data were collected using pClamp 10 and Clampex 10.4 software (Molecular Devices, Palo Alto, CA), sampled at 10 kHz and filtered at 3 kHz, analyzed with Clampfit 10.4.

### Rotarod test

Rotarod test was performed using a five lane rotarod treadmill (Ugo Basile). Mice were trained for 5 consecutive days, one trial per day. One trial lasted for 300 s while the rotarod was accelerated from 4 to 40 rpm, or from 8 to 80 rpm. Mice were carefully placed on each lane of the rotarod and acclimated for 30 s. Experimenter checked the latency to fall of each mouse. Latency was regarded as 300 s when a mouse withstood the full 300 s.

### Immunofluorescence

For brain sectioning, mice were perfusion fixed. The brains were postfixed using 4% paraformaldehyde (PFA) (wj0012, Genesion), cryoprotected in 30% sucrose, and sectioned into 6 μm coronal sections. Coronal sections were sliced throughout the brain. Immunofluorescence was performed on 6 μm thick paraffin or frozen sections. The paraffin sections of brain were deparaffinized, rehydrated, and permeabilized in 10 mM sodium citrate buffer. After being restored to room temperature, the frozen sections of brain were first soaked in PBS for 10 min and then blocked for 30 min. Then, all sections were incubated with primary antibodies overnight at 4 °C. After 5 times washing with PBS, the sections were incubated with Alexa Fluor 488 and 568 conjugated secondary antibodies (Life Technologies). Finally, the sections were imaged using a Zeiss LSM 900 or a Zeiss LSM 710 with a 10× objective after incubation with DAPI solution (D9542, Sigma–Aldrich, USA) for 10 min.

For culture cells, cells were fixed with 4% paraformaldehyde (PFA) (pH 7.4) (Genesion, wj0012) for 30 min at room temperature. Then the cells were incubated with primary antibodies in PBS containing 10% goat serum, 0.3% Triton X-100 (Solarbio, T8200) overnight at 4°C, washed 3 times with PBS, and then incubated with secondary antibodies for 1 h. Fluorescence images were acquired on Leica or Zeiss confocal microscope. 4′,6-diamidino-2-phenylindole (DAPI) (Sigma, D9542) was used to label nuclei. The primary antibodies were mouse anti-DNA antibody (Abcam, ab27156, 1:200), rabbit anti-TFAM antibody (Abcam, ab176558 1:1000), rabbit anti-TOM20 antibody (Abcam, ab186735, 1:200), polyclonal anti-SYN1 antibody (rabbit, 1:200, Sigma-Aldrich,574777), monoclonal anti-NeuN (rabbit, 1:1000, Abcam, ab177487), monoclonal anti-GFAP (mouse, 1:500, Sigma-Aldrich, MAB360). Alexa Fluor 488 (Thermo Fisher Scientific, A-11001, 1:500), Alexa Fluor 568, (Thermo Fisher Scientific, A-11011, 1:500).

### Cell culture and neural differentiation

Normal human iPSCs 90D, as we previously reported,[32] were cultured in mTesR1 medium (Stem Cell, 85850). Colonies, following 0.5 mM EDTA (Aladdin, E116428) treatment, were cultured on Matrigel (Becton Dickinson, 356234) coated plates. All cultures were maintained in a humidified incubator containing 5% CO_2_ at 37°C. Neurons were differentiated from 90D iPSCs. Briefly, 0.5 mM EDTA-dissociated iPSCs were split 1:4 on Matrigel-coated plates. On the following day, the iPSC medium was replaced with N2B27 medium, i.e.1:1 mixture of DMEM/F12 (Hyclone, SH30023-018) and neurobasal medium (Gibco, 21103-049) supplemented with 0.5×N2 (Gibco, 17502048), 0.5×B-27 (Gibco, 17504044), 1×Glutamax (Gibco, 35050-061),1×NEAA (Gibco, 11140-050) and 1×penicillin/streptomycin (Gibco, SV30010). 3 μM CHIR99021 (Selleck, S1263), 2 μM Dorsomorphin (Selleck, S7840) and 2 μM SB431542 (Selleck, S1067) were added in the medium. The culture medium was changed every other day. 90D iPSCs were maintained under the above condition for 7 days, followed by dissociation with 1 mg/ml Dispase (Gibco, 17105041) and passage at 1:6 in the medium described above. Then 1 μM CHIR99021, 2 μM Dorsomorphin, 2 μM SB431542 were added for another 7 days for differentiation into neural progenitor cells (NPCs). The NPCs were expanded with N2B27 medium containing 20 ng/ml EGF (Sino Biological, 10605-HNAE-1) and 20 ng/ml bFGF (Sino Biological, 10014-HNAE-1) and passaged with accutase (Sigma, A6964). For neuron differentiation, NPCs were plated onto matrigel-coated plates and cultured in N2B27 medium supplemented with 10 ng/mL BDNF (BioVision, 4004) and 1 μM cAMP (Sigma, A6885).

HEK 293T (ATCC, YS-ATCC207) were grown in DMEM-Dulbecco’s modified Eagles medium, High Glucose (Hyclone, SH30022-2B) supplemented with 10% fetal bovine serum (Natocor, SFBE) and penicillin/streptomycin. Trypsin-EDTA (0.25T) (Gibco, 25200056)-dissociated 293T were split 1:3 on 10 cm plates.

### Plasmid construct and Lentivirus production, infection

The plasmid pDsRed2-mito (Clontech, 632421), with a mitochondrial targeting sequence from COX8A (cytochrome c oxidase subunit 8A) fused to the fluorescence protein DsRed2, was used to mark mitochondria. Mito-GFP was constructed by replacing DsRed2 with GFP. ER-DsRed plasmid was kindly provided by Prof. György Hajnóczky (Thomas Jefferson University, Philadelphia, PA, USA). Split-GFP (spGFP1-10-ERt and Mitot-2×spGFP11) by Prof. Chao Tong (Zhejiang University, China). UL12.5 by Prof. Tao Peng (Guangzhou Institutes of Biomedicine and Health, CAS, China). Mito-DsRed, UL12.5-DsRed, mito-GFP, ER-DsRed, split-GFP plasmids were subcloned into a lentiviral expression vector—pRlenti. UL12.5 plasmids were also subcloned into a lentiviral expression vector—pSin-puro.

HEK 293T cells were plated onto 10 cm dishes and co-transfected with target plasmids and packaging vectors (PMD2.G (Addgene, 12259) and PSPAX2 (Addgene, 12260) for pRlenti and pSin vector) using polyethylenimine (PEI) (Sigma, 765090), Opti-MEM I Reduced Serum Medium(Thermo Fisher, 31985-070) - based transfection. The viruses were filtered with 0.45-μm filters (Millipore, SLHV033RB) and centrifuged at 50,000 g for 2.5 h. The precipitant was used to infect cells after 10-14 days of neural differentiation without polybrene. pSin vector infected cells followed by selection with 1 μg/ml puromycin (Selleck, S7417) for 3-4 days.

### Quantitative reverse transcription PCR (qRT-PCR)

Total RNA was extracted from neurons with TRIzol (Invitrogen, 15596026) 1 week after infection, and 2 μg of RNA was used to generate complementary DNA. For mitochondrial genomic level detection, total DNA was extracted from neurons using a Universal Genomic DNA Kit (TIANGEN, DP304) 1 week after infection and diluted to a concentration of 3 ng/mL. qPCR experiments were performed with SYBR Green Master Mix (Takara, RR420L) and primers specific for *MTND1, MTND4, MTATP8, ATP5A, MIRO1, MIRO2, MILT1, MILT2, ITPR3*, *RYR1*, *SIRT1*, *PPARGC1A*, *MTOR*, *PRKAB1*, *PRKAA2*, *mtNd1, mtNd4, mtCox1, mtCox3, mtCytb,* and UL12.5 using a Bio-Rad® CF96 Real-Time PCR System. Thermal condition was 95°C for 10 min, followed by 40 cycles of 95°C for 15 s, 60°C for 15 s and 72°C for 15 s. The expression was normalized using GAPDH as an internal reference. Primer sequences are provided in Supplementary Table 1.

### Western blot analyses

The neuron cells after 1 week of infection were lysed in RIPA buffer (Beyotime, P0013K) containing both protease inhibitor cocktail (Bimake, B14002) and PMSF (Beyotime, ST506) and subjected to Western blot analysis. Total proteins, electrophoresed on 12-15% polyacrylamide gel containing sodium dodecyl sulfate, were immediately transferred onto the 0.45 μM PVDF membranes (Millipore, ISEQ00010). Membranes were blocked in 5% milk, followed by incubation for 1 h at room temperature or overnight at 4°C with primary antibodies. Then membranes were washed in TBST, incubated with secondary antibodies for 1 h at room temperature, and washed in TBST. ECL substrate solution (Millipore, WBKLS0500) was added for chemiluminescence, the intensity of which was quantitated using a chemiluminescence imager.

The primary antibodies were Rabbit polyclonal anti ATP5A (Proteintech, 14676-1-AP, 1:1000), Rabbit polyclonal anti MTATP8 (Santa Cruz, sc-84231, 1:1000), Rabbit polyclonal to Tubulin β-3 (TUBB3) (Biolegend, 802001, 1:1000) and Rabbit polyclonal to GAPDH (Abcam, ab9485, 1:1000), Rabbit polyclonal anti MTCOX3 (Proteintech, 55082-1-AP,1:500), Mouse monoclonal anti FLAG (Sigma-Aldrich, F1804, 1:1000).

### OCR and ECAR measurements using Seahorse Cellular Flux assays

Seahorse plates were coated with 0.1 mg/ml Poly-D-lysine and matrigel thereafter. NPC were passaged and seeded in N2B27 medium supplemented with 10 ng/mL BDNF and 1 μM cAMP for neural differentiation onto pre-treated Seahorse plates with 5 × 10^5^ cells per XF24 well to ensure about 90% surface coverage at the time of the experiment. After 3 days differentiation, mito-DsRed or UL12.5-DsRed virus were used to infect neurons, media were exchanged as above. After infection for 2 weeks, for ECAR analysis, medium was exchanged for glycolysis stress medium (i.e. XF Base medium (Agilent, 102353) supplemented with 2 mM glutamine) at 1 h before the assay. The cultures were then incubated in a non-CO_2_ incubator at 37°C to equilibrate. Substrates and selective inhibitors were injected during the measurement to achieve final concentrations of glucose at 2.5 mM, oligomycin at 1 μM, and 2-DG at 100 mM, according to the Seahorse XF Glycolysis Stress Test Kit (Agilent, 103020-100) manufacturer’s instructions. For OCR analysis, medium was exchanged for mitochondrial stress medium (i.e. XF Base medium supplemented with 2 mM glutamine and 10 mM glucose) at 1 h before the assay. The cultures were similarly equilibrated. Substrates and selective inhibitors were injected to achieve final concentrations of oligomycin at 1 μM, FCCP at 0.5 μM and rotenone/antimycin A at 1.0 µM, according to the Seahorse XF Cell Mito Stress Test Kit (Agilent, 103015-100) manufacturer’s instructions. The OCR and ECAR values were further normalized to the number of cells present in each well. The baselines of OCR and ECAR were defined as the average values. Changes in OCR and ECAR in response to substrates and inhibitors were defined as the maximal change after the chemical addition as compared with the baseline.

### Electrophysiology

Whole-cell patch clamp recordings were performed from cells infected with mito-DsRed or UL12.5-DsRed virus after 2 weeks of differentiation. The bath was constantly perfused with fresh HEPES-buffered saline. The recording micropipettes (tip resistance 3–6 MU) were filled with internal solution. Recordings were made using Axopatch 200B amplifier (Axon Instruments). Signals were filtered at 2 kHz and sampled at 5 kHz. The whole-cell capacitance was fully compensated. The series resistance was uncompensated but monitored during the experiment by the amplitude of the capacitive current in response to a 10 mV pulse. All recordings were performed at room temperature and chemicals were purchased from Sigma. Frequency and amplitude of spontaneous postsynaptic currents were measured with the Mini Analysis Program software (Synaptosoft, Leonia, NJ).

### Assessment of beading and fragmented neurites

The method for assessment of beading and fragmented neurites was described in our previous report.[32] Briefly, mito-DsRed/UL12.5-DsRed-labeled neurons were treated with 2-DG for 2 or 4 days. For the calcium inhibitors treatment, 1 mM EGTA (Sigma, E4378), 1 µM BAPTA-AM (Abcam, ab120503), 10 µM Dan (Abcam, ab142274), 2 µM XesC (Abcam, ab120914), 2-APB (Selleck, S6657), and Ryanodine (Abcam, ab120083) were added simultaneously with 2-DG for 2 days. Neurons were observed under a phase-contrast microscope. More than 100 neurons were assessed blindly in three independent trials. The fraction of neurites with neuritic beads was calculated as a percentage of total neurites.

### Measurement of mitochondrial motility

Mito-DsRed/UL12.5-DsRed-labeled neurons were transferred to Nunc™ Glass Bottom Dishes (Thermo Scientific™, 150680) and treated with 2-DG for 12 h, and then mitochondrial movements were recorded for 10 min at 4/s using Zeiss confocal microscope maintained at 37 °C. Multiple Kymograph measurement was analyzed as in our previous report.[20] Mitochondria were subsequently classified as motile (velocity > 0.1 μm/s) or stationary (velocity < 0.1 μm/s).

### **ΔΨ**_m_ measurement

For measuring ΔΨ_m_, neurons were treated with or without 1 μg/mL oligomycin (Cell Signaling Technology, 9996L) for 30 min, then loaded with 50 nM Dilc(5) (Molecular Probes, M34151) and oligomycin for another 30 min in 37°C, and then washed twice with serum-free medium before imaging.

### Flow cytometry analysis

Standard flow cytometry analysis was performed with BD Fortessa. Mitochondria-ER contact were analyzed using a genetically encoded reporter split super-folder GFP protein as previously reported.[21] In brief, mito-DsRed/UL12.5-DsRed-labeled neurons were infected with split-GFP virus. The cells harvested at day 4, were spun at 500 g for 5 min and the resuspend pellet was used for flow analysis.

### Assessment of ER fragmentation and profile analysis

To assess neuritic ER fragmentation, ER-DsRed was used to mark neuritic ER. Confocal microscopy was used to detect the morphology of ER. For analyzing the physical contacts of ER and mitochondria, ER and mitochondria were marked by ER-DsRed and mito-GFP viruses, respectively. The profiles of ER and mitochondria were analyzed by ImageJ software.

### Intracellular calcium detection

Fluo-4 AM (Invitrogen, F-14201) was used to measure intracellular calcium. Cells were incubated with 1 µM Fluo-4 AM for 30 min at 37°C, and then washed twice with medium before imaging. The cells were detected by laser confocal or fluorescence microscopy with the excitation wavelength of 494 nm and emission wavelength of 516 nm.

### Mitochondrial calcium detection

Mito-GCaMP5 was used to measure mitochondrial calcium. Cells expressing mito-DsRed/UL12.5-DsRed were infected with mito-GCaMP5, and then treated with 2-DG for 24 h, washed twice with medium before imaging. The cells were detected by confocal microscopy with the excitation wavelength of 494 nm and emission wavelength of 516 nm.

### Live cell calcium imaging

Mito-GCaMP5 was used to measure mitochondrial calcium and ER-XCaMPRed was used to measure ER calcium. Cells expressing FLAG/UL12.5-3FLAG were infected with mito-GCaMP5 and ER-XCaMPRed, and then treated with 2-DG for 24 h, washed twice with medium before imaging. The cells were detected by laser confocal microscopy with wavelength of 488 nm and 561 nm.

### RNA-seq analysis

Total RNA was extracted with TRIzol. Libraries were prepared using an Illumina TruSeq RNA Sample Prep kit (Illumina, 20020594) following the manufacturer’s instruction. The RNA-seq experiment was performed at Annoroad Gene Technology Co. Ltd. (Beijing, China). The raw reads of RNA-seq were trimmed from 3′ to 5′ ends for each read, and the reads shorter than 50 bp were discarded. The reads that passed the quality control were mapped to mouse genome (mm10) with STAR (version 2.7). Stringtie (version 2.1.4) were then used to estimate transcript abundance in fragments per kilobase per million (FPKM). Genes were annotated as differentially expressed using DESeq2 (version 1.44). The GO analysis and KEGG pathway analysis were performed using R packages.

### MTT assay

Cell proliferation was determined by the MTT assay according to the instructions of a MTT Cell Proliferation and Cytotoxicity Assay Kit (Beyotime Biotechnology). Briefly, NPCs were passaged and seeded in N2B27 medium supplemented with 10 ng/mL BDNF and 1 μM cAMP for neural differentiation onto 24-well culture plates with 5 × 10 ^4^ NPCs per well. After 3 days differentiation, mito-DsRed or UL12.5-DsRed virus were used to infect neurons, media were exchanged as above. After infection for 2 weeks, for MTT assay, the neurons were incubated with MTT solution for 4 hr. 150 μL formazan solvent was added and shaken for 5 min until the crystals were dissolved. The optical density (OD) was determined with a microplate reader (Bio-Rad 550) at 570 nm wavelength.

### Statistical analysis

Data were plotted in GraphPad Prism (Version 8; La Jolla, CA, USA) software. Statistical analysis was performed using unpaired student’s t test or one-way ANOVA.

## Supporting information

Supple FigureLegends

## Acknowledgements

We thank all the members in the lab of Prof. Xingguo Liu. This work is funded by the National Key Research and Development Program of China (2022YFE0210100, 2019YFA0904500, 2023YFE0210100, 2022YFA1103800), the National Natural Science Foundation projects of China (32025010, 92157202, 32241002, 92254301, 92357302, 32261160376, 31970709, 32070729, 32100619, 32170747, 32322022, 32370782, 32371007, 32300608, 32300620), NSFC/RGC Joint Grant Scheme 2022/2023 (N_CUHK 428/22), the Strategic Priority Research Program of the Chinese Academy of Sciences (XDB0480000), the Key Research Program, CAS (ZDBS-ZRKJZ-TLC003), International Cooperation Program, CAS (154144KYSB20200006), CAS Project for Young Scientists in Basic Research (YSBR-075), Guangdong Province Science and Technology Program (2024A1515030120, 2023B0303000023, 2023B1111050005, 2023A1515030231, 2022A1515110493, 2023B1212060050, 2021A1515012513, 2021B1515020096, 2022A1515012616, 2022A1515110951, 2023B1212120009, 2024A1515010782, 2024B1515040020), Guangzhou Science and Technology Program (202102021037, 202102020827, 202102080066, 202206060002, 2023A04J0414), Health@InnoHK funding support from the Innovation Technology Commission of the Hong Kong SAR, Basic Research Project of Guangzhou Institutes of Biomedicine and Health, Chinese Academy of Sciences and CAS Youth Innovation Promotion Association (to Y. W and K. C).

## Author Contributions

L. Z., F. B. and J. Z. Contributed equally to this work. X. L. initiated and supervised the project. L. Z. and F. B. designed and performed the experiments. G. C., J. Z., Y. Q., L. H. and H. W. participated in electrophysiological experiments. Y.D. analyzed RNA-seq. M. L., J. X., Q. L., and Q. M. participated in the experiment. Y. W. and L. Y. participated in the manuscript revision. X. L., F. B., L. Z., and G. C. wrote the manuscript.

## References

[1] A. Lim, R. H. Thomas, European journal of paediatric neurology: EJPN: official journal of the European Paediatric Neurology Society 2020, 24, 47.

[2] G. Zsurka, W. S. Kunz, The Lancet. Neurology 2015, 14, 956.

[3] T. Iizuka, F. Sakai, N. Suzuki, T. Hata, S. Tsukahara, M. Fukuda, Y. Takiyama, Neurology 2002, 59, 816.

[4] D. M. Kirby, K. J. Rennie, T. K. Smulders-Srinivasan, R. Acin-Perez, M. Whittington, J. A. Enriquez, A. J. Trevelyan, D. M. Turnbull, R. N. Lightowlers, Cell proliferation 2009, 42, 413.

[5] a) M. T. Todorova, P. Tandon, R. A. Madore, C. E. Stafstrom, T. N. Seyfried, Epilepsia 2000, 41, 933; b) A. E. Greene, M. T. Todorova, R. McGowan, T. N. Seyfried, Epilepsia 2001, 42, 1371.

[6] a) W. Duan, Z. Guo, H. Jiang, M. Ware, X. J. Li, M. P. Mattson, Proceedings of the National Academy of Sciences of the United States of America 2003, 100, 2911; b) N. Maswood, J. Young, E. Tilmont, Z. Zhang, D. M. Gash, G. A. Gerhardt, R. Grondin, G. S. Roth, J. Mattison, M. A. Lane, R. E. Carson, R. M. Cohen, P. R. Mouton, C. Quigley, M. P. Mattson, D. K. Ingram, Proceedings of the National Academy of Sciences of the United States of America 2004, 101, 18171.

[7] a) S. Someya, G. C. Kujoth, M. J. Kim, T. A. Hacker, M. Vermulst, R. Weindruch, T. A. Prolla, PLoS One 2017, 12, e0171159; b) S. Santra, R. W. Gilkerson, M. Davidson, E. A. Schon, Ann Neurol 2004, 56, 662.

[8] a) E. Volmering, P. Niehusmann, V. Peeva, A. Grote, G. Zsurka, J. Altmüller, P. Nürnberg, A. Becker, S. Schoch, C. Elger, W. Kunz, Acta neuropathologica 2016, 132, 277; b) C. Perier, A. Bender, E. García-Arumí, M. Melià, J. Bové, C. Laub, T. Klopstock, M. Elstner, R. Mounsey, P. Teismann, T. Prolla, A. Andreu, M. Vila, Brain: a journal of neurology 2013, 136, 2369.

[9] A. Spinazzola, F. Invernizzi, F. Carrara, E. Lamantea, A. Donati, M. Dirocco, I. Giordano, M. Meznaric-Petrusa, E. Baruffini, I. Ferrero, M. Zeviani, Journal of inherited metabolic disease 2009, 32, 143.

[10] a) R. M. Epand, R. F. Epand, B. Berno, L. Pelosi, G. Brandolin, Biochemistry 2009, 48, 12358; b) P. E. Bonnen, J. W. Yarham, A. Besse, P. Wu, E. A. Faqeih, A. M. Al-Asmari, M. A. Saleh, W. Eyaid, A. Hadeel, L. He, F. Smith, S. Yau, E. M. Simcox, S. Miwa, T. Donti, K. K. Abu-Amero, L. J. Wong, W. J. Craigen, B. H. Graham, K. L. Scott, R. McFarland, R. W. Taylor, American journal of human genetics 2013, 93, 471; c) H. Chen, M. Vermulst, Y. E. Wang, A. Chomyn, T. A. Prolla, J. M. McCaffery, D. C. Chan, Cell 2010, 141, 280.

[11] P. Verstreken, C. V. Ly, K. J. T. Venken, T.-W. Koh, Y. Zhou, H. J. Bellen, Neuron 2005, 47, 365.

[12] A. Bender, K. J. Krishnan, C. M. Morris, G. A. Taylor, A. K. Reeve, R. H. Perry, E. Jaros, J. S. Hersheson, J. Betts, T. Klopstock, R. W. Taylor, D. M. Turnbull, Nat. Genet. 2006, 38, 515.

[13] a) B. A. Duguay, J. R. Smiley, Journal of virology 2013, 87, 11787; b) L. Yang, Q. Long, J. Liu, H. Tang, Y. Li, F. Bao, D. Qin, D. Pei, X. Liu, Cellular and molecular life sciences: CMLS 2015, 72, 2585.

[14] a) D. Goertsen, N. C. Flytzanis, N. Goeden, M. R. Chuapoco, A. Cummins, Y. Chen, Y. Fan, Q. Zhang, J. Sharma, Y. Duan, L. Wang, G. Feng, Y. Chen, N. Y. Ip, J. Pickel, V. Gradinaru, Nature neuroscience 2022, 25, 106; b) A. Konno, H. Hirai, Journal of neuroscience methods 2020, 346, 108914.

[15] a) R. K. Naviaux, W. L. Nyhan, B. A. Barshop, J. Poulton, D. Markusic, N. C. Karpinski, R. H. Haas, Ann Neurol 1999, 45, 54; b) A. H. Hakonen, P. Isohanni, A. Paetau, R. Herva, A. Suomalainen, T. Lönnqvist, Brain: a journal of neurology 2007, 130, 3032.

[16] J. Liu, C. Reeves, Z. Michalak, A. Coppola, B. Diehl, S. M. Sisodiya, M. Thom, Acta Neuropathol Commun 2014, 2, 71.

[17] S. Vielhaber, D. Kunz, K. Winkler, F. R. Wiedemann, E. Kirches, H. Feistner, H. J. Heinze, C. E. Elger, W. Schubert, W. S. Kunz, Brain: a journal of neurology 2000, 123, 1339.

[18] J. R. Gledhill, M. G. Montgomery, A. G. Leslie, J. E. Walker, Proceedings of the National Academy of Sciences of the United States of America 2007, 104, 15671.

[19] P. Gonzalez-Rodriguez, E. Zampese, K. A. Stout, J. N. Guzman, E. Ilijic, B. Yang, T. Tkatch, M. A. Stavarache, D. L. Wokosin, L. Gao, M. G. Kaplitt, J. Lopez-Barneo, P. T. Schumacker, D. J. Surmeier, Nature 2021.

[20] F. X. Bao, H. Y. Shi, Q. Long, L. Yang, Y. Wu, Z. F. Ying, D. J. Qin, J. Zhang, Y. P. Guo, H. M. Li, X. G. Liu, CNS Neurosci Ther 2016, 22, 648.

[21] Z. Yang, X. Zhao, J. Xu, W. Shang, C. Tong, J Cell Sci 2018, 131.

[22] R. Villegas, N. W. Martinez, J. Lillo, P. Pihan, D. Hernandez, J. L. Twiss, F. A. Court, J Neurosci 2014, 34, 7179.

[23] I. Amigo, S. L. Menezes-Filho, L. A. Luevano-Martinez, B. Chausse, A. J. Kowaltowski, Aging Cell 2017, 16, 73.

[24] a) E. Salińska, J. W. Lazarewicz, Neurologia i neurochirurgia polska 1996, 30 Suppl 2, 35; b) T. Yamamoto, A. Takahara, Current topics in medicinal chemistry 2009, 9, 377.

[25] T. Hayashi, T. P. Su, Cell 2007, 131, 596.

[26] a) C. Vicidomini, L. Ponzoni, D. Lim, M. J. Schmeisser, D. Reim, N. Morello, D. Orellana, A. Tozzi, V. Durante, P. Scalmani, M. Mantegazza, A. A. Genazzani, M. Giustetto, M. Sala, P. Calabresi, T. M. Boeckers, C. Sala, C. Verpelli, Molecular psychiatry 2017, 22, 689; b) B. C. Orem, A. Rajaee, D. P. Stirling, Journal of neurotrauma 2022, 39, 311.

[27] J. Schmiedel, S. Jackson, J. Schafer, H. Reichmann, J. Neurol. 2003, 250, 267.

[28] C. Costa, V. Belcastro, A. Tozzi, M. Di Filippo, M. Tantucci, S. Siliquini, A. Autuori, B. Picconi, M. G. Spillantini, E. Fedele, A. Pittaluga, M. Raiteri, P. Calabresi, J. Neurosci. 2008, 28, 8040.

[29] a) G. Zsurka, W. S. Kunz, J. Bioenerg. Biomembr. 2010, 42, 443; b) J. M. Shoffner, M. T. Lott, A. M. Lezza, P. Seibel, S. W. Ballinger, D. C. Wallace, Cell 1990, 61, 931.

[30] M. Cherubini, L. Lopez-Molina, S. Gines, Neurobiol Dis 2020, 136, 104741.

[31] E. L. Wilson, E. Metzakopian, Cell Death Differ. 2021, 28, 1804.

[32] F. Bao, H. Shi, M. Gao, L. Yang, L. Zhou, Q. Zhao, Y. Wu, K. Chen, G. Xiang, Q. Long, J. Guo, J. Zhang, X. Liu, Cell death & disease 2018, 9, 966.

[33] a) L. V. Vandervore, R. Schot, C. Milanese, D. J. Smits, E. Kasteleijn, A. E. Fry, D. T. Pilz, S. Brock, E. Börklü-Yücel, M. Post, N. Bahi-Buisson, M. J. Sánchez-Soler, M. van Slegtenhorst, B. Keren, A. Afenjar, S. A. Coury, W. H. Tan, R. Oegema, L. S. de Vries, K. A. Fawcett, P. G. J. Nikkels, A. Bertoli-Avella, A. Al Hashem, A. A. Alwabel, K. Tlili-Graiess, S. Efthymiou, F. Zafar, N. Rana, F. Bibi, H. Houlden, R. Maroofian, R. E. Person, A. Crunk, J. M. Savatt, L. Turner, M. Doosti, E. G. Karimiani, N. W. Saadi, J. Akhondian, M. H. Lequin, H. Kayserili, P. J. van der Spek, A. C. Jansen, J. M. Kros, R. M. Verdijk, N. J. Milošević, M. Fornerod, P. G. Mastroberardino, G. M. S. Mancini, American journal of human genetics 2019, 105, 1126; b) S. B. Wortmann, F. M. Vaz, T. Gardeitchik, L. E. Vissers, G. H. Renkema, J. H. Schuurs-Hoeijmakers, W. Kulik, M. Lammens, C. Christin, L. A. Kluijtmans, R. J. Rodenburg, L. G. Nijtmans, A. Grünewald, C. Klein, J. M. Gerhold, T. Kozicz, P. M. van Hasselt, M. Harakalova, W. Kloosterman, I. Barić, E. Pronicka, S. K. Ucar, K. Naess, K. K. Singhal, Z. Krumina, C. Gilissen, H. van Bokhoven, J. A. Veltman, J. A. Smeitink, D. J. Lefeber, J. N. Spelbrink, R. A. Wevers, E. Morava, A. P. de Brouwer, Nat Genet 2012, 44, 797.

[34] a) F. Xiao, J. Zhang, C. Zhang, W. An, Laboratory investigation; a journal of technical methods and pathology 2017, 97, 289; b) V. Basso, E. Marchesan, C. Peggion, J. Chakraborty, S. von Stockum, M. Giacomello, D. Ottolini, V. Debattisti, F. Caicci, E. Tasca, V. Pegoraro, C. Angelini, A. Antonini, A. Bertoli, M. Brini, E. Ziviani, Pharmacological research 2018, 138, 43; c) D. Lau, N. Hartopp, N. Welsh, S. Mueller, E. Glennon, G. Mórotz, A. Annibali, P. Gomez-Suaga, R. Stoica, S. Paillusson, C. Miller, Cell death & disease 2018, 9, 327.

[35] a) P. E. Coskun, M. F. Beal, D. C. Wallace, Proc. Natl. Acad. Sci. U. S. A. 2004, 101, 10726; b) A. Esteves, A. Domingues, I. Ferreira, C. Januário, R. Swerdlow, C. Oliveira, S. Cardoso, Mitochondrion 2008, 8, 219.

[36] a) D. Zala, M. V. Hinckelmann, H. Yu, M. M. Lyra da Cunha, G. Liot, F. P. Cordelières, S. Marco, F. Saudou, Cell 2013, 152, 479; b) M. V. Hinckelmann, A. Virlogeux, C. Niehage, C. Poujol, D. Choquet, B. Hoflack, D. Zala, F. Saudou, Nature communications 2016, 7, 13233.

[37] a) B. R. Desousa, K. K. Kim, A. E. Jones, A. B. Ball, W. Y. Hsieh, P. Swain, D. H. Morrow, A. J. Brownstein, D. A. Ferrick, O. S. Shirihai, A. Neilson, D. A. Nathanson, G. W. Rogers, B. P. Dranka, A. N. Murphy, C. Affourtit, S. J. Bensinger, L. Stiles, N. Romero, A. S. Divakaruni, EMBO reports 2023, 24, e56380; b) A. I. Tarasov, E. J. Griffiths, G. A. Rutter, Cell Calcium 2012, 52, 28.

[38] S. Li, G. J. Xiong, N. Huang, Z. H. Sheng, Nat Metab 2020, 2, 1077.

[39] X. Wang, T. L. Schwarz, Cell 2009, 136, 163.

