## Supplementary material for "Calorie Restriction Induces Degeneration of Neurons with Mitochondrial DNA Depletion by Altering ER-Mitochondria Calcium Transfer": Supple FigureLegends

**Supplementary Figure Legends**

**Figure S1** Expression of UL12.5 reduces mtRNA in mice brain.

A) Quantification of RNA levels of UL12.5 in mice brain and liver by qPCR. Error bars: SD (n=3; ***P < 0.001, * p < 0.05).
B) Western blot analysis of UL12.5 expression in liver.
C) Quantification of brain RNA levels of mitochondrial encoded proteins (*mtNd1, mtNd4 mtCox1, mtCox3, mtCytb*) by qPCR. Error bars: SD (n=3; ***P < 0.001).

**Figure S2** I(K) peak amplitude and excitatory-to-inhibitory(E/I) in UL12.5 mouse hippocampi.

A) The evoked action potential (APs) amplitude peaks in the hippocampus neurons from control and UL12.5 mice. Error bars: SD (n =13 neurons from 3 mice; *P <0.05)
B) The peak amplitude of voltage-activated potassium channel currents (I(K)) in the hippocampus from control and UL12.5 mice. Error bars: SEM (n =13 neurons from 3 mice).
C) The frequency of spontaneous inhibitory postsynaptic current (sIPSC) in hippocampal excitatory neurons from control and UL12.5 mice. Error bars: SD (n = 10 neurons from 3 mice).
D) The amplitude of sIPSC in hippocampal excitatory neurons from control and UL12.5 mice. Error bars: SD (n =10 neurons from 3 mice).
E) The excitatory-to-inhibitory(E/I) amplitudes in hippocampal excitatory neurons from control and UL12.5 mice. Error bars: SD (n = 12 neurons from 3 mice for control, n = 7 neurons from 3 mice for UL12.5).

**Figure S3** Transcriptome analysis in UL12.5 mice.

A) Expression profiles (heat map) of mtDNA- and nuclear-encoded mitochondrial transcripts of UL12.5 and control mice (red, high gene expression; blue, low gene expression).

B, C) Gene Ontology (GO) and Kyoto Encyclopedia of Genes and Genomes (KEGG) pathway analysis of the differentially expressed genes in UL12.5 mice. B) Significant GOs. C) Significant KEGG signaling pathways.

**Figure S4** Expression of UL12.5 disrupts mitochondrial nucleoid structure.

A) Representative images of a fixed UL12.5-DsRed (red) expressing neuron with mitochondrial marker protein TOM20 (green). Scale bar: 10 μm.
B) Mitochondrial nucleoid was labeled by anti-TFAM antibody in control and UL12.5 expressing neurons. Representative confocal microscopy images are shown. Scale bar: 10 μm.
C) qPCR was used to detect RNA levels of mitochondrial encoded (*MTND1, MTND4 MTATP8*) and nuclear encoded *ATP5A*. Error bars: SEM (n=4; ***P < 0.001).

**Figure S5** Glycolysis capacity in rho^-^ neurons.

A) Quantification of the OCR for each treatment. Error bars: SEM (n≥3; ***P < 0.001).

B, C) Neurons were infected with mito-DsRed or UL12.5-DsRed before ECAR analysis, followed by sequential addition of glucose, oligomycin (which inhibits ATP synthesis), and 2-DG (which inhibits glycolysis) (B). Quantification of the ECAR for each cellular stressor is shown in (C). Error bars: SEM (n=3; **P < 0.01).

D) Glycolysis (subtraction of the basal ECAR from the glucose ECAR), glycolytic capacity (subtraction of the basal ECAR from the oligomycin ECAR), and glycolytic reserve (subtraction of the glucose ECAR from the oligomycin ECAR) are shown. Error bars: SEM (n=3; *P < 0.01).

**Figure S6** The electrophysiological activity of rho^-^ neurons.

A) Plots show the average current–voltage relationships in control and rho^-^ neurons. Recordings were made in voltage-clamp mode (holding potential l −70 mV) and currents were elicited by a series of voltage steps from −50 to +60 mV. Error bars: SEM (n = 11).

B,C) Plots show normalized current–voltage relationships of sodium channels (B) and potassium channels (C) from control and rho^-^ neurons. The peak current amplitudes were normalized to 1. Error bars: SEM (n ≥ 18; * p < 0.05; ***, p < 0.001).

D, E) Activating inward currents induced by 500 μM glutamate in control and rho^-^ neurons (D), quantification of the peak current amplitudes indicates that there is no significant different in rho^-^ neurons (E). Error bars: SEM (n≥16).

F, G) Activating inward currents induced by 500 μM GABA in control and rho^-^ neurons (F) and quantification of the peak current amplitudes (G). Error bars: SEM (n≥15).

H) Action potentials recorded after injection of current steps (−50 to 90 pA) from control and rho^-^ neurons.

**Figure S7** CR by 2-DG, rapamycin or low glucose reduces cell viability.

A-C) Quantification of the expression of calorie restriction related genes (SIRT1, PPARGC1A, mTOR, PRKAB1, PRKAA2) in control or UL12.5 expressing neurons treated with 2DG (A), rapamycin (B), or low glucose (C) by qPCR. Error bars: SD (n=3; ***P < 0.001, **P < 0.01, *P < 0.05).

D-F) Cell viability was assessed by manual cell count from control and rho^-^ neurons with or without 2-DG, rapamycin, or low glucose treatment for 4 days. Error bars: SD (n=3; *P < 0.05).

**Figure S8** 2-DG has no effect on mitochondrial motility in rho^-^ neurons.

A, B) Mitochondrial kymographs were recorded from control and rho^-^ neurons with or without 2-DG treatment for 12 h. The percentages of anterograde (A) and retrograde (B) moving mitochondria are shown. Error bars: SEM (n≥22 mitochondria).

**Figure S9** Contacts of mitochondria and ER increase in rho^-^ neurons. 2-DG induces intracellular calcium increases in rho^-^ neurons.

A) Representative images of split-GFP in control and rho^-^ neurons. Scale bar: 10 μm.

B) Representative images of mito-GFP- and ER-DsRed-expressing rho^-^ neurons. Profile analysis of the white lines is shown in the lower panels. Scale bar: 10 μm.

C, D) Control and rho^-^ neurons were treated with 2-DG for 2 days, followed by staining with Fluo4. Representative confocal microscopy images are shown in (C). Scale bar:10 μm. Quantification of the Fluo4 fluorescence intensities (D). Intensities are normalized to the mean of the control neurons, Error bars: SEM (n=28 neurons from three biological replicates, *** P < 0.001).

E-G) Representative images of mito-GCaMP5- and ER-XCaMPRed-expressing rho^-^ neurons. Scale bar: 10 μm. Quantification of the mito-GCaMP5 fluorescence intensities (F), and quantification of the ER-XCaMPRed fluorescent intensity (G), Error bars: SD (n≥60 neurons from three biological replicates, ** P < 0.01, *** P < 0.001).

**Figure S10** Transcriptome analysis reveals calcium signaling pathway dysfunction in rho^-^ neurons.

A, B) Gene Ontology (GO) and Kyoto Encyclopedia of Genes and Genomes (KEGG) pathway analysis of the differentially expressed genes in rho^-^ neurons. A) Significant GOs. B) Significant KEGG signaling pathways.

C) Representative calcium signaling pathways constructed by KEGG. Red and green genes represent up- and down-regulated genes in rho^-^ neurons, respectively.

**Figure S11** Neuritic degeneration induced by 2-DG in rho^-^ neurons was prevented by 2-APB or Ryanodine.

Images of control and rho^-^ neurons after 2-DG with or without simultaneous 2-APB or Ryanodine treatment for 2 days. Scale bar: 10 μm.

**Figure S12** ShRNA mediated knock-down of IP3R1 and RYR2 mRNA in neurons.

A) Expression of *IP3R3* and *RYR1* mRNA in control and rho^-^ neurons by RT-qPCR. Error bars: SD (n=2, * P < 0.05, ** P < 0.01).

B) Efficiency of shRNA-mediated knock-down of *IP3R3* and *RYR1* mRNA was measured by RT-qPCR. Error bars: SD (n=3, *** P < 0.001).
