## Supplementary figures and images for "Calorie Restriction Induces Degeneration of Neurons with Mitochondrial DNA Depletion by Altering ER-Mitochondria Calcium Transfer"

### Figure S1

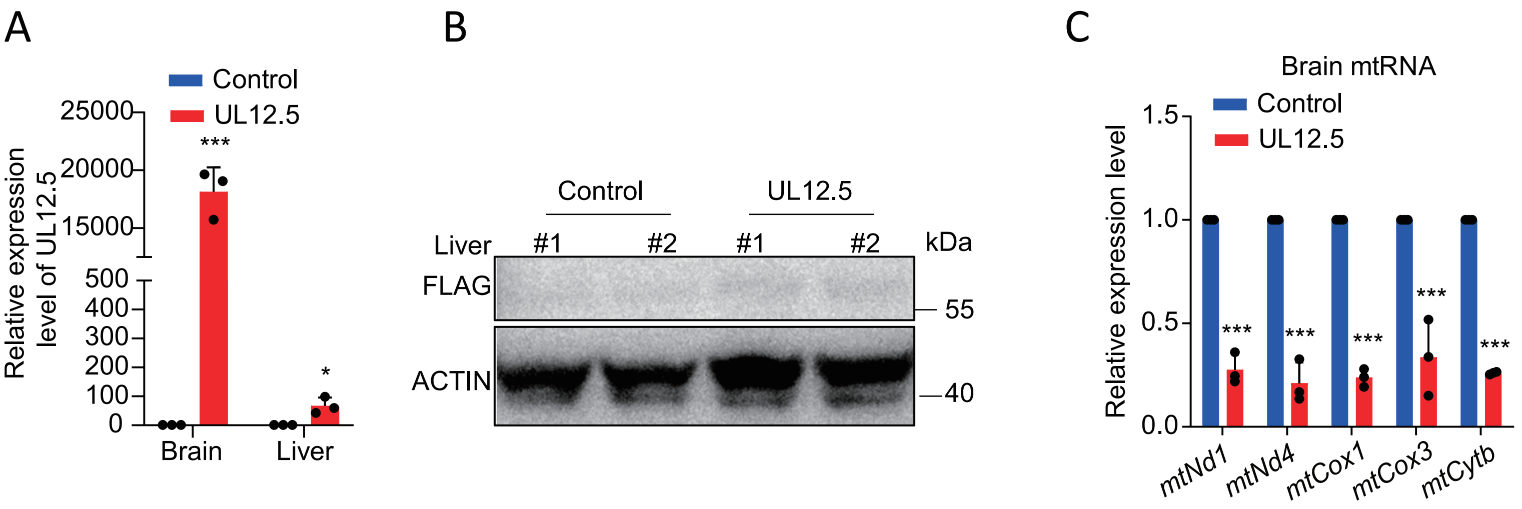

### Figure S2

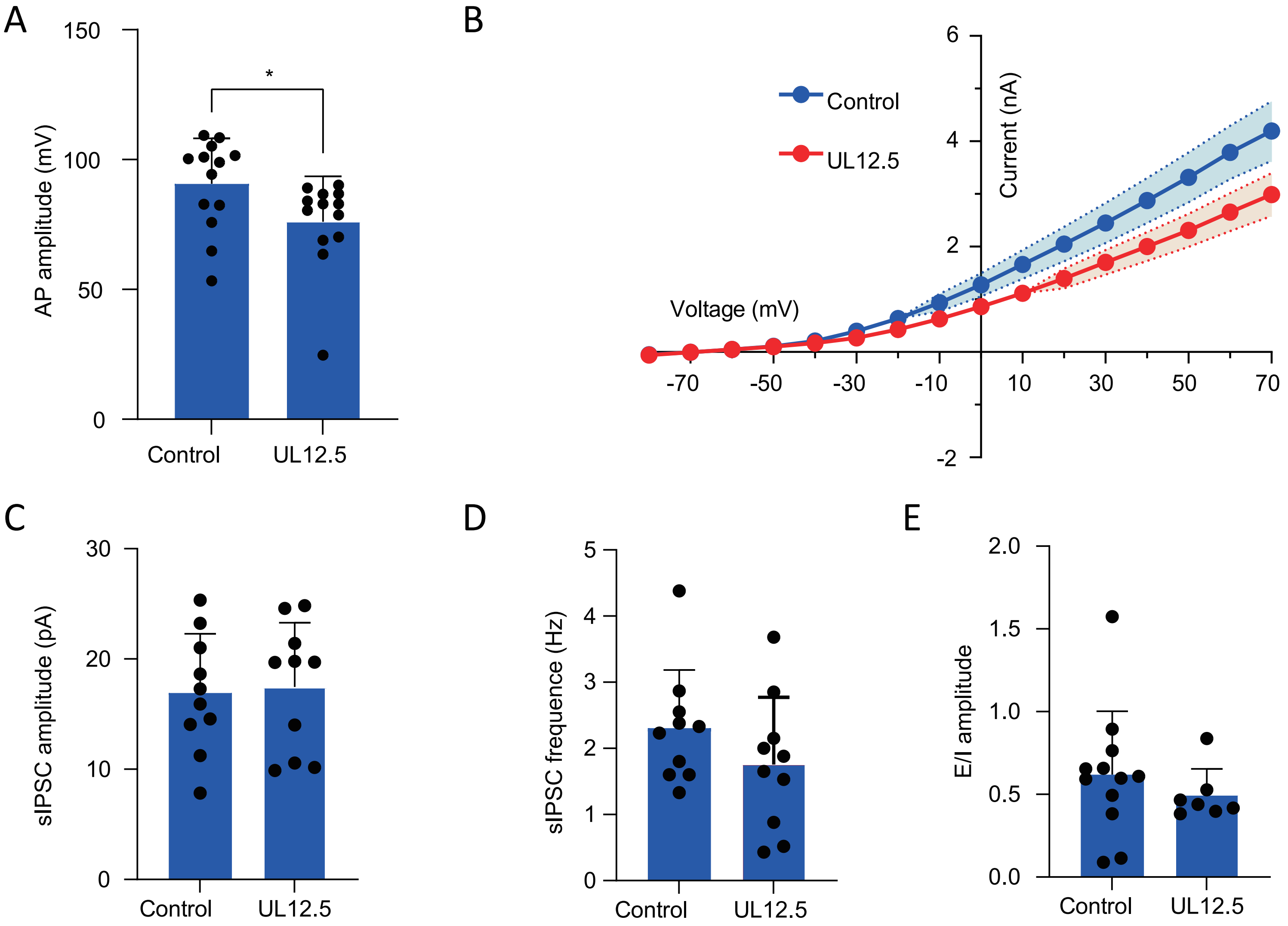

### Figure S3

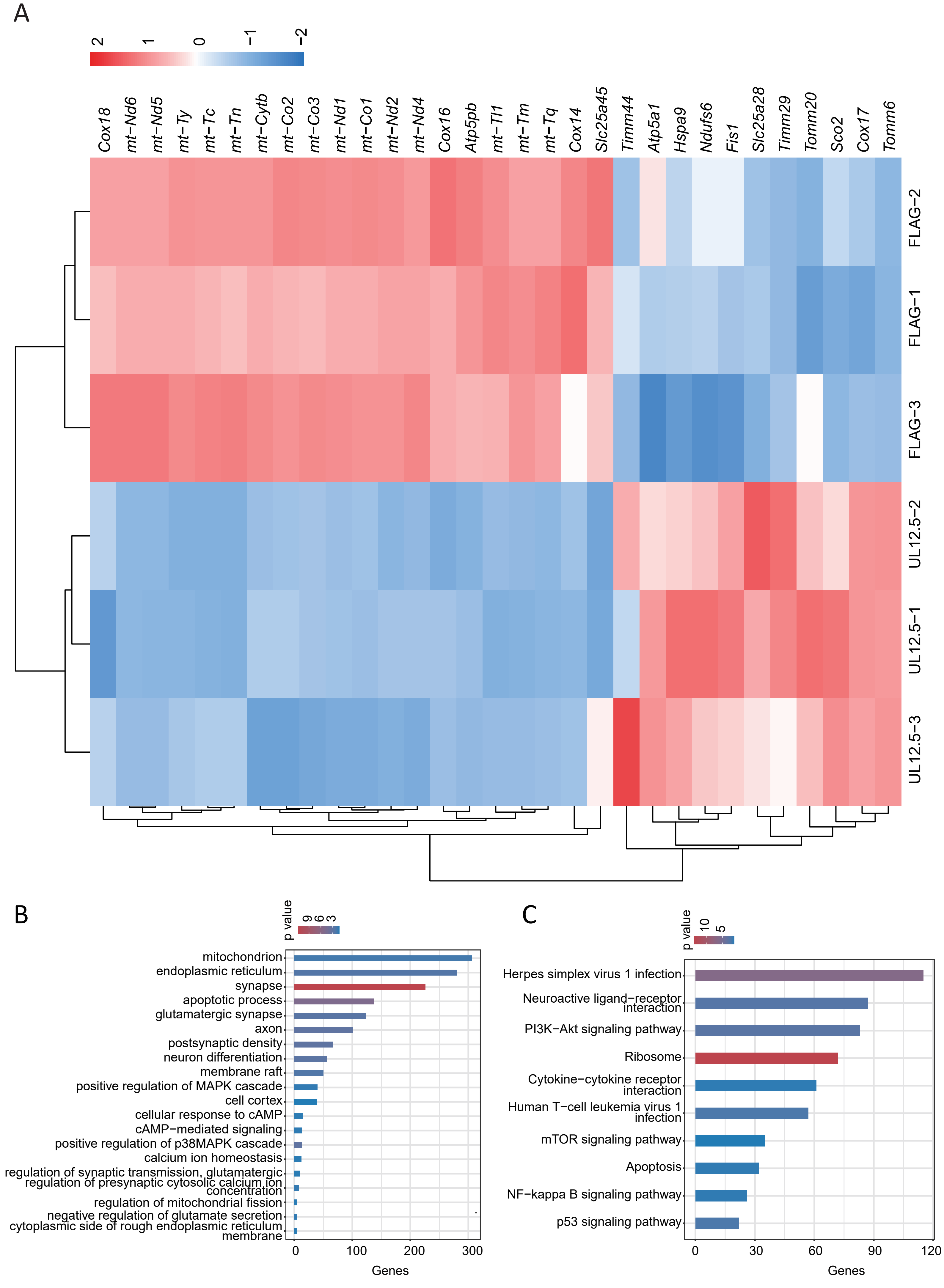

### Figure S4

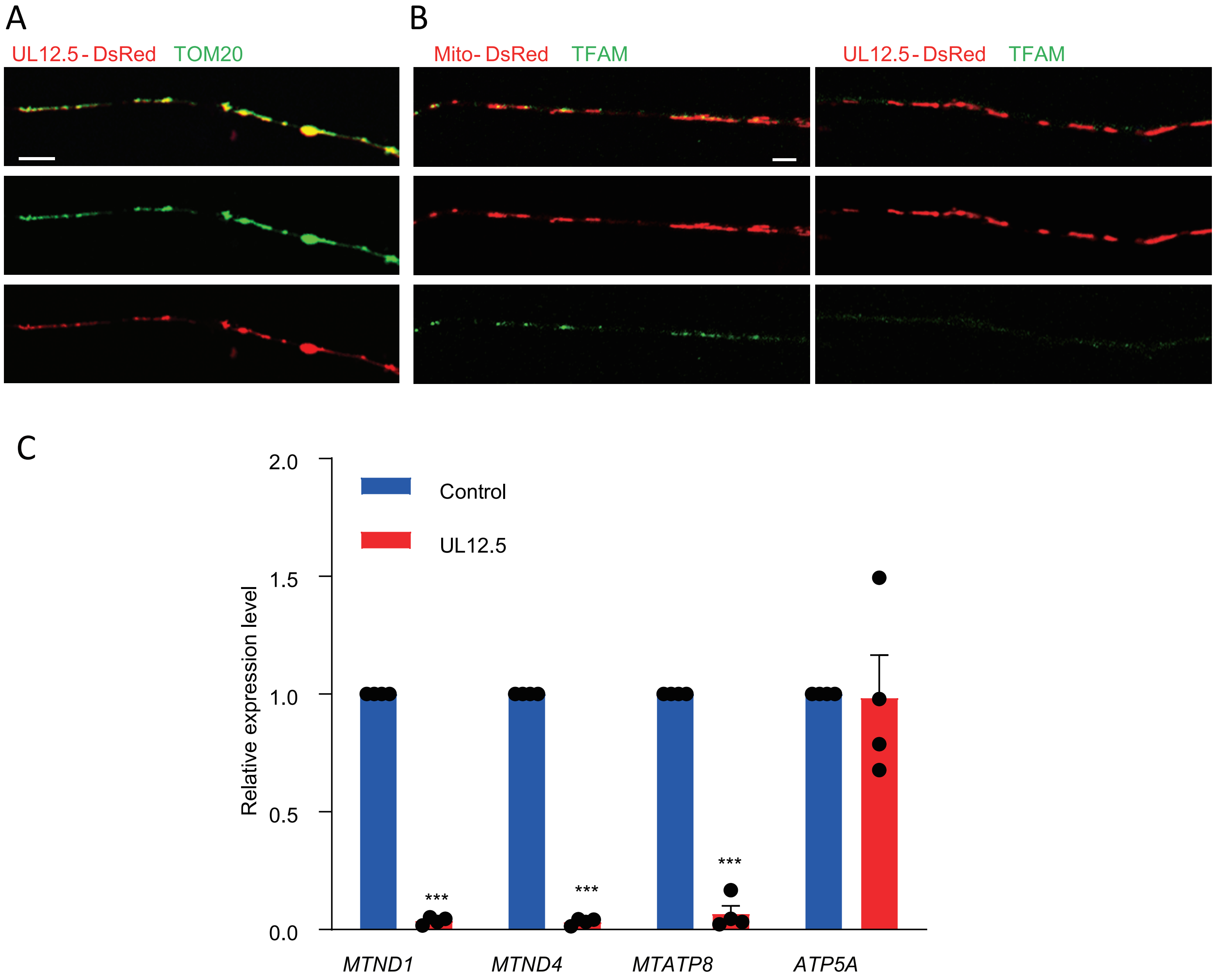

### Figure S5

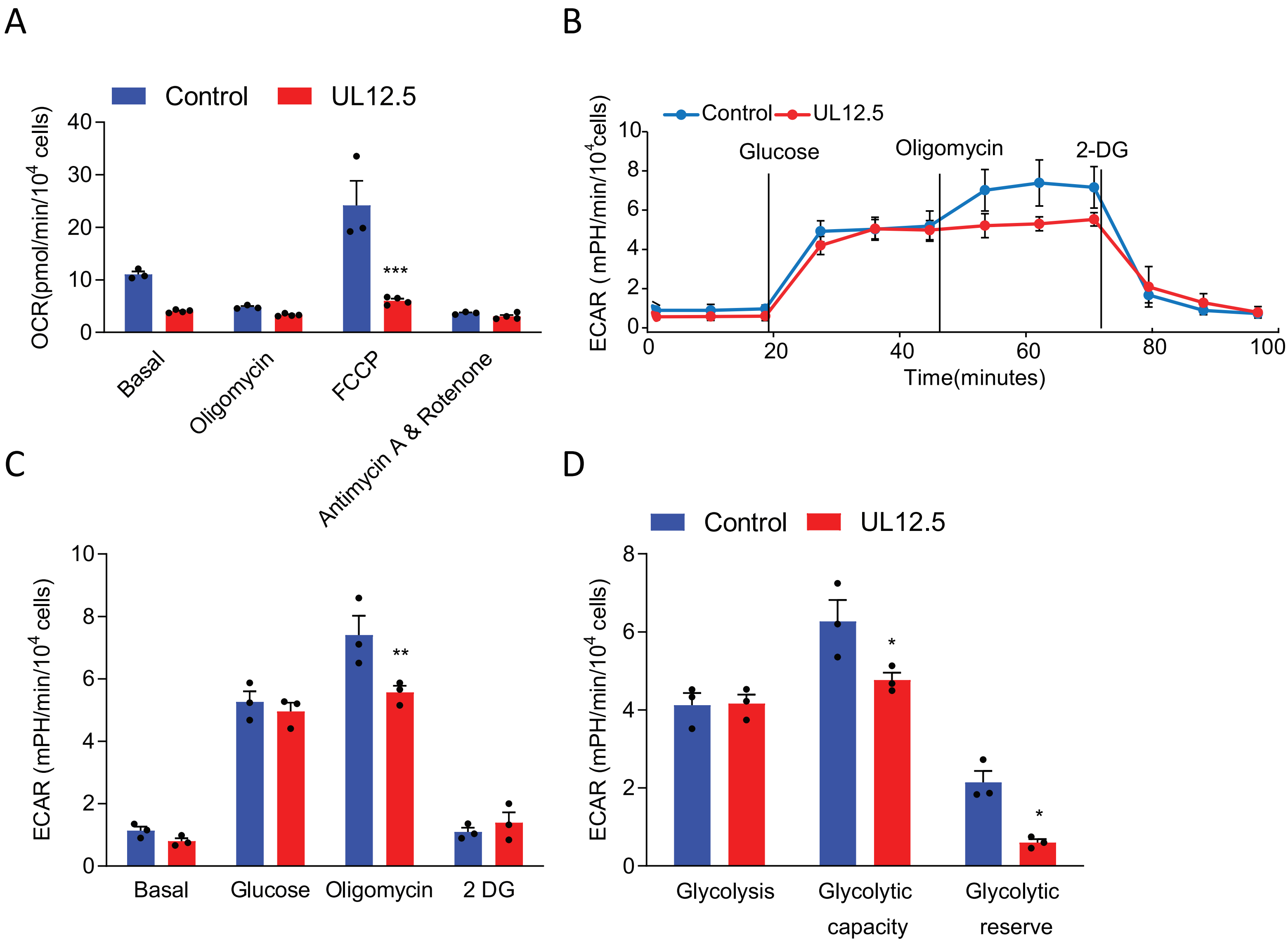

### Figure S6

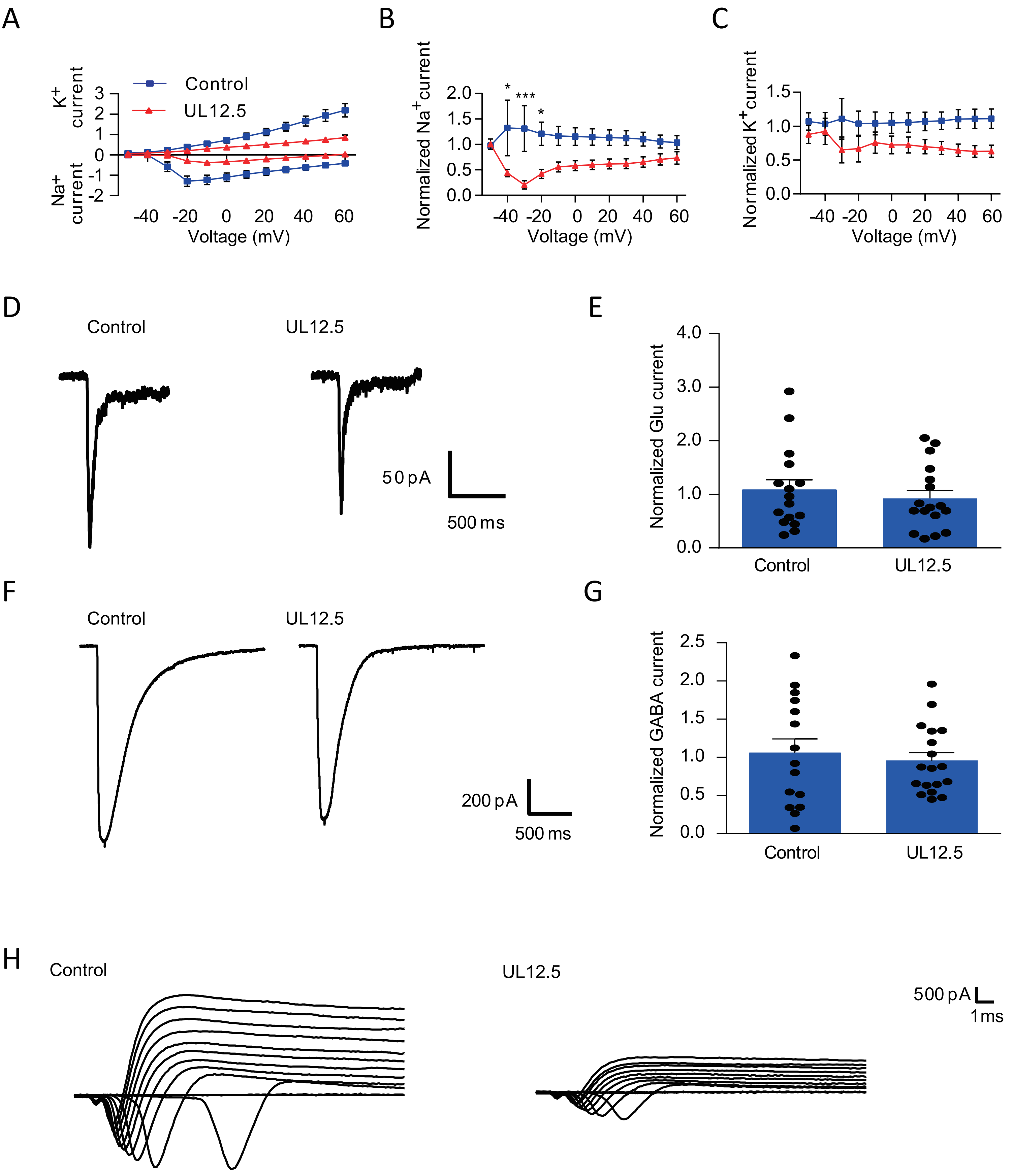

### Figure S7

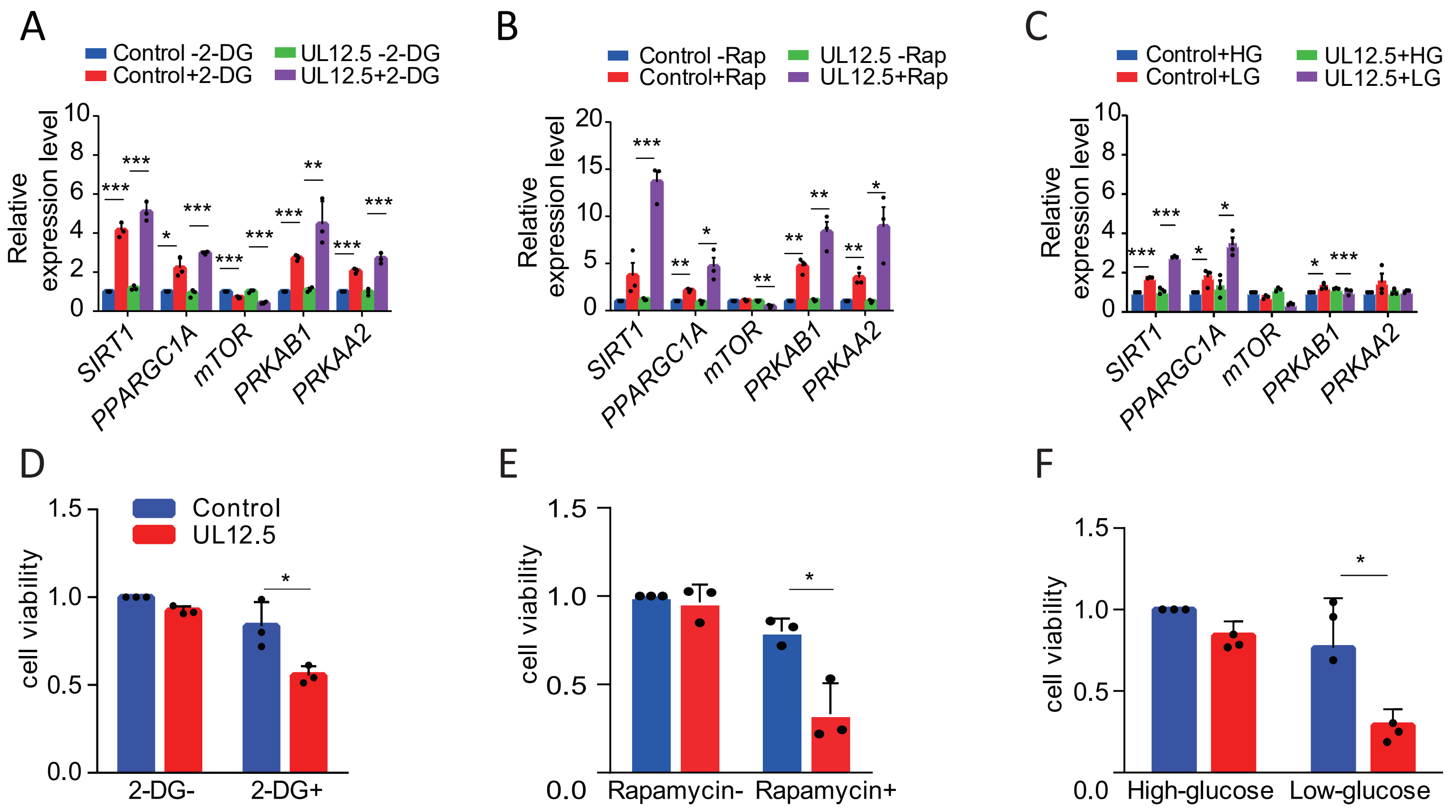

### Figure S8

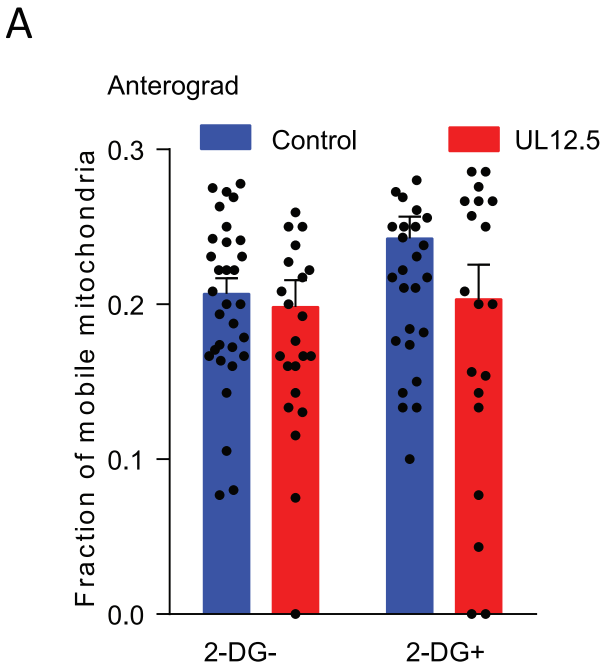

### Figure S9

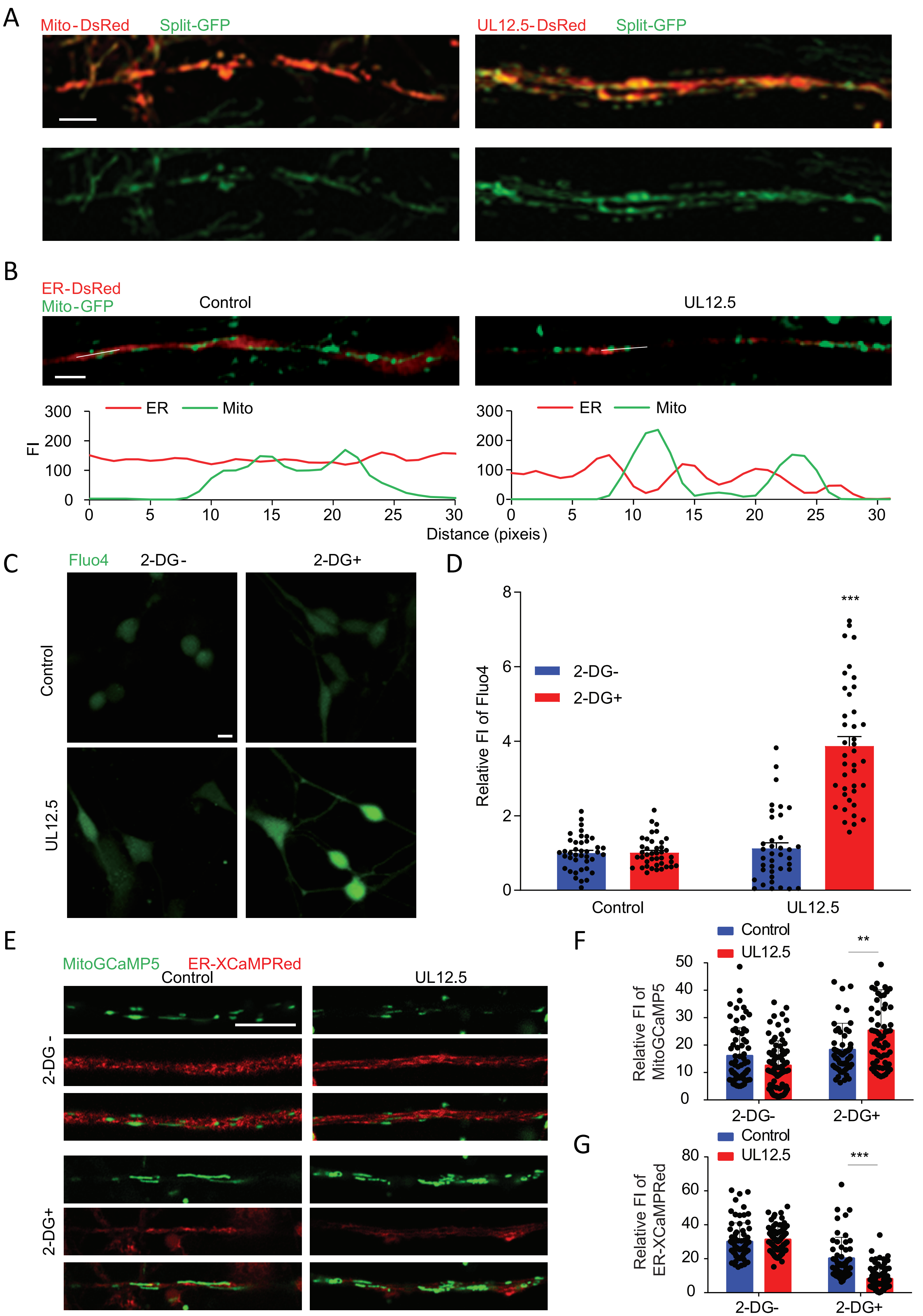

### Figure S10

#### Figure S10

A

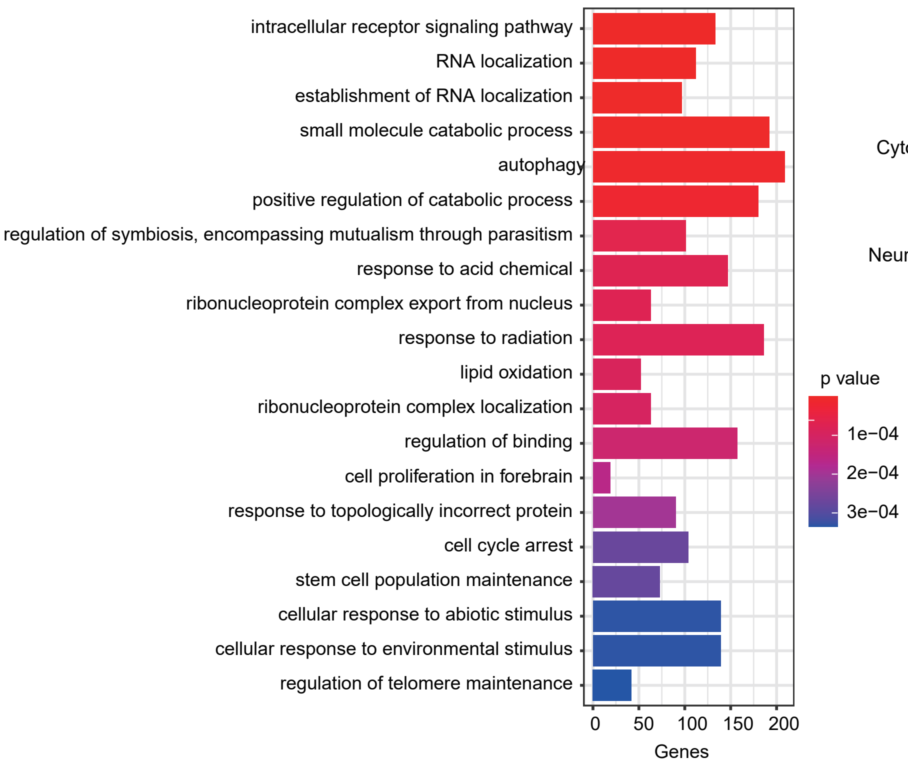


B

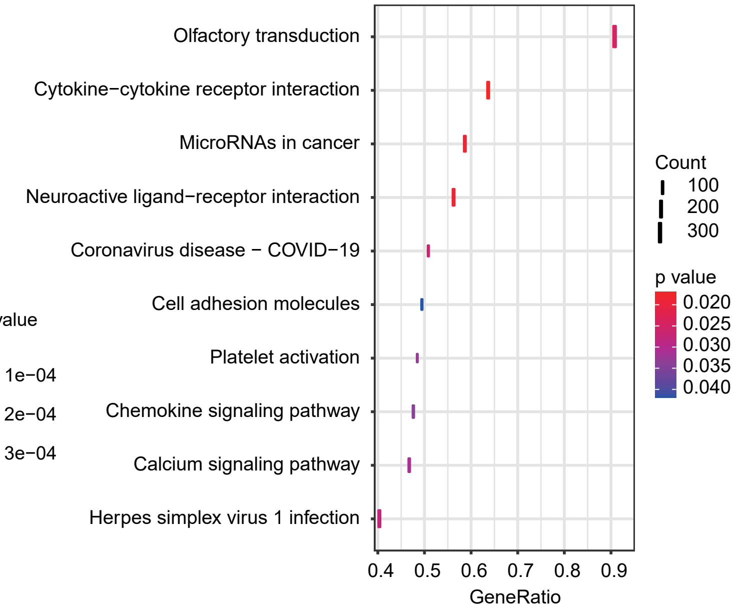


C

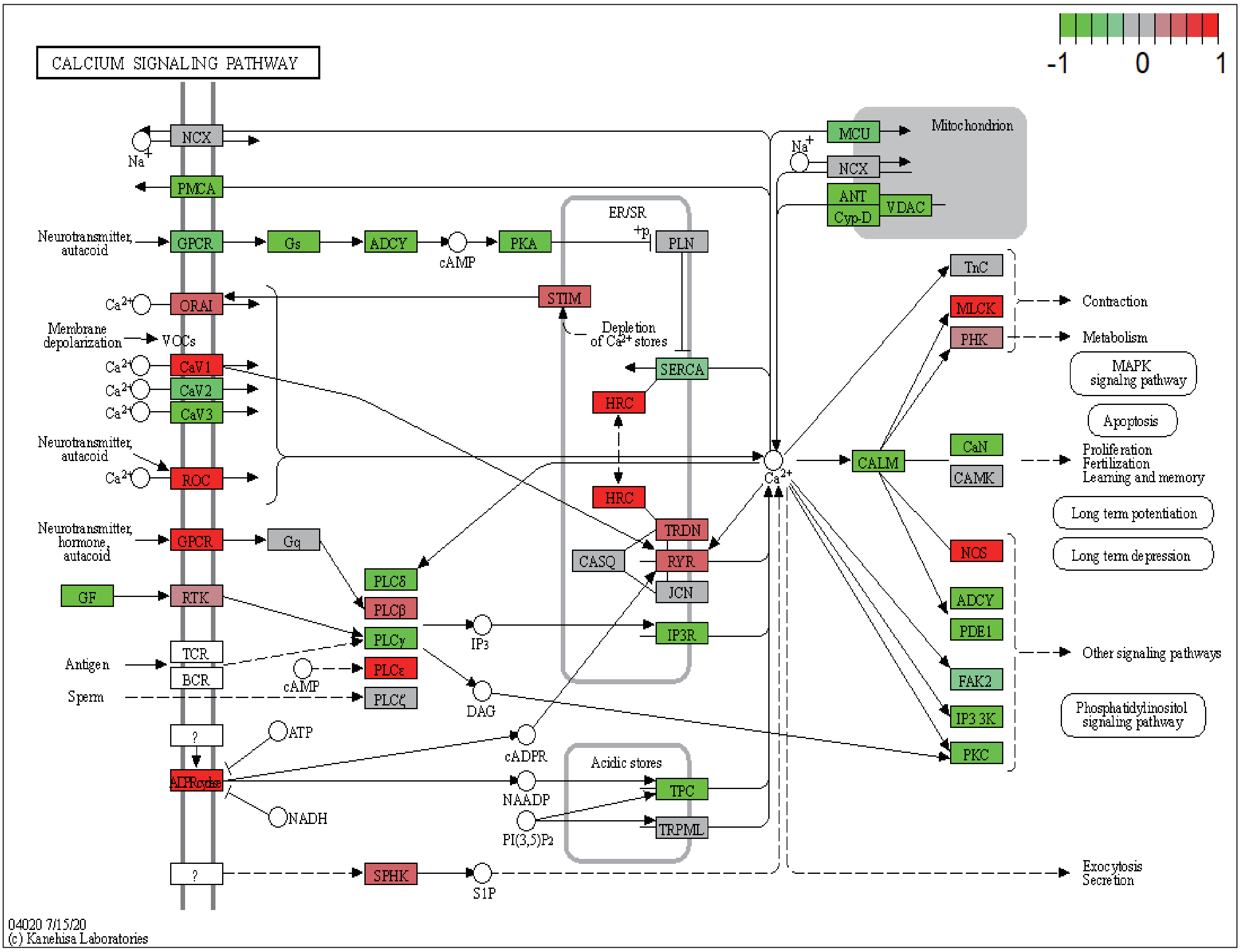

### Figure S11

#### Figure S11

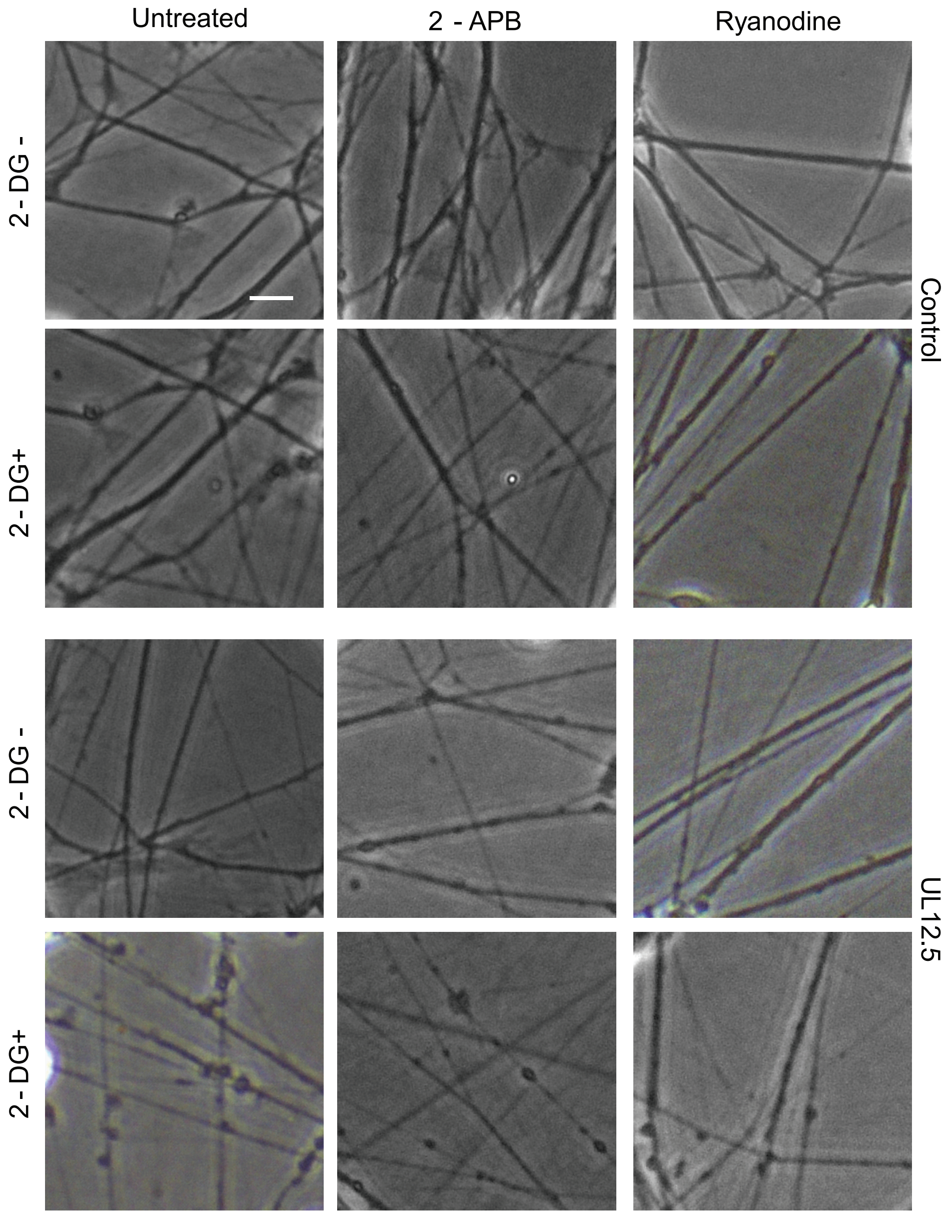

### Figure S12

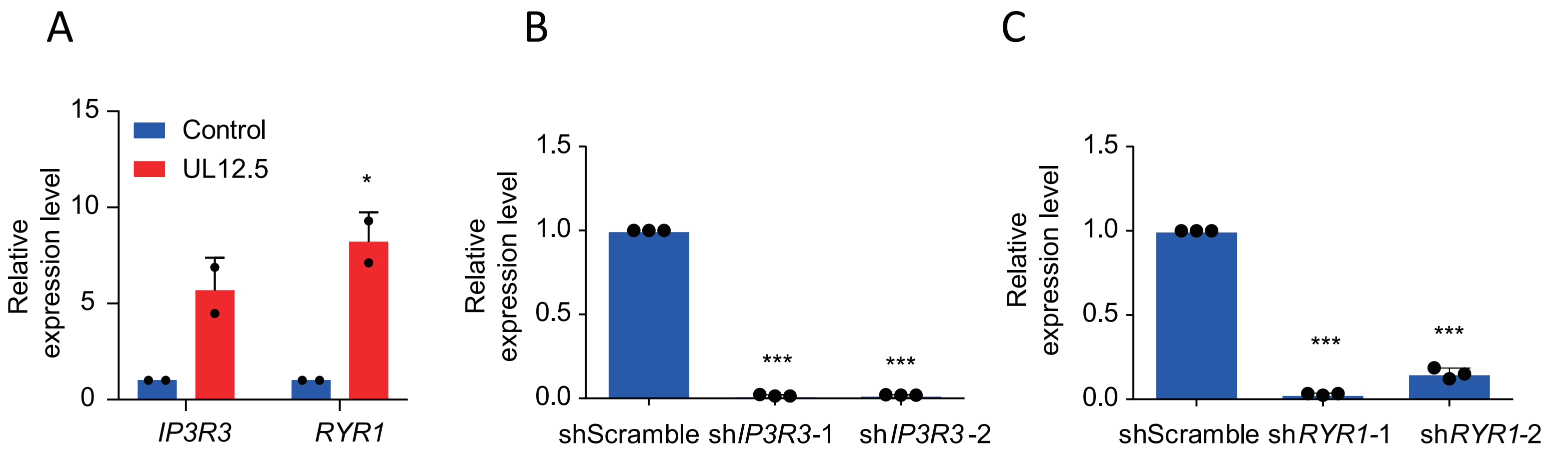
